# IL-21-STAT3 axis negatively regulates LAIR1 expression in B cells

**DOI:** 10.1101/2025.01.14.632971

**Authors:** Kenji Yoshida, Izumi Kurata-Sato, Yemil Atisha-Fregoso, Cynthia Aranow, Betty Diamond

**Affiliations:** Center of Autoimmune Musculoskeletal and Hematopoietic Diseases, The Feinstein Institutes for Medical Research, Northwell Health, Manhasset, New York, USA; Laboratories for Pharmacology, Pharmaceuticals Research Center, Asahi Kasei Pharma Corporation, Shizuoka, Japan; Japan Drug Development and Medical Affairs, Eli Lilly Japan K.K., Kobe, Hyogo, Japan

## Abstract

LAIR1 is an inhibitory receptor broadly expressed on human immune cells, including B cells. LAIR1 has been shown to modulate BCR signaling, however, it is still unclear whether its suppressive activity can be a negative regulator for autoreactivity. In this study, we demonstrate the LAIR1 expression profile on human B cells and prove its regulatory function and relationships to B cell autoreactivity. We show that both the frequency and level of LAIR1 expression decreases during B cell differentiation. LAIR1 expressing (LAIR1^+^) switched memory (SWM) B cells have a transcriptional profile less differentiated toward a plasma cell (PC) phenotype, harbor more autoreactive B cells and exhibit less PC differentiation *in vitro* than the LAIR1 negative (LAIR1^-^) counterpart. These data suggests that LAIR1 functions as a B cell tolerance checkpoint. We confirm previous data showing that patients with systemic lupus erythematosus (SLE) express less LAIR1 on B cells, implying a breakdown of the checkpoint, consistent with the enhanced PC differentiation seen in SLE. We further demonstrate that LAIR1 expression is down-regulated through the IL-21/STAT3 pathway which is known to be upregulated in SLE. These data suggest therapeutic targets that might decrease the aberrant PC differentiation observed in SLE.

## Introduction

Systemic lupus erythematosus (SLE) is an autoimmune disease with dysregulation of B cells, and ensuring autoantibody production. Emerging data suggest a dysregulation of memory B cells and overexuberant plasma cell (PC) differentiation as well as hyperactivity of naïve B cells (1, 2).

In healthy individuals, autoreactive B cells are strictly limited by several tolerance checkpoints to avoid undesirable activation and differentiation to PC (3, 4). Autoreactive B cells are not effectively controlled in patients with SLE and secretion of disease-specific autoantibodies and immune complex lead to inflammation and organ damage, such as lupus nephritis (5–7).

LAIR1 is an inhibitory receptor with immunoreceptor tyrosine-based inhibitory motifs (ITIMs) in its intracellular domain (8). Activation of LAIR1 by its ligands, collagen (9) and C1q (10), induces phosphorylation of the ITIMs followed by recruitment of the phosphatase SHP-1 (11–13), which regulates kinase cascades. In human B cells, LAIR1 has been shown to suppress B cell receptor (BCR)-induced Ca^2+^ influx and downstream kinase cascades when it is brought into proximity with the BCR (14–16). Colombo et al investigated LAIR1 expression on CD20^+^ IgM^+^ B cells in patients with SLE and found that the number of B cells not expressing LAIR1 (LAIR1^-^) was expanded compared to healthy donors (15). However, the role of LAIR1 throughout B cell differentiation and the mechanisms that regulate its expression are still unknown. Here we show that LAIR1^+^ switched memory (SWM) B cells have a transcriptional profile less differentiated toward a PC phenotype, contain more autoreactive cells, and exhibit less PC differentiation after stimulation than LAIR1^-^ SWM B cells. *In vitro culture* of B cells showed loss of LAIR1 expression following activation; it is mediated by IL-21/STAT3 pathway. These observations provide a plausible explanation for the reduced LAIR1 on B cells in SLE and suggests therapeutic strategies that might maintain LAIR1 expression as a checkpoint on B cell activation and PC differentiation.

## Materials and methods

### Reagents

All reagents in this study are summarized in Supplementary Table 1, 2, 3 and 4.

### Preparation of PBMCs

PBMCs from healthy donors (HD) were obtained by Ficoll-Pague gradient centrifugation of leukopaks purchased from the New York Blood Center. PBMCs were similarly obtained from blood of patients with SLE collected in heparinized tubes. The patients were recruited at the Feinstein Institutes for Medical Research. Characteristics of the patients and healthy donors are summarized in Supplementary Table 5.

### Flow cytometry

For phenotyping by flow cytometry, cells were stained for 20 minutes with antibodies against cell membrane proteins and Fc blocker (Miltenyi Biotec), except for the staining with anti-IgG, followed by washing with 2% FBS in HBSS. Live cells were identified on the basis of Fixable Viability Dye eFluor 780 (Thermo Fisher Scientific) labeling. The gating strategy for B cell subsets is shown in Supplementary Figures (Supplementary Figure. 1A, 1B, 1C, 3A, and 3B).

For ANA staining for autoreactive BCR, nuclear extract from HeLa cells was used as previously described (3). Incubation with nuclear extract was performed in 1.5% nonfat dry milk (LabScientific) in HBSS. After thorough washing, cells were stained with streptavidin-APC (Biolegend) and antibodies against B cell markers. The gating strategy for ANA^+^ B cells is shown in Supplementary Figures (Supplementary Figure. 1D, 3F and 3G).

Cells were analyzed with a BD Symphony instrument or sorted with a BD FACS Aria. The data were further analyzed with Flowjo software (v10.9.0).

### Bulk RNA sequencing

B cells isolated with EasySep Human B Cell Isolation Kit (Stem cell technologies) were stained with the antibody cocktail; PerCP-Cy5.5 anti-CD19 (1:100), APC anti-CD27 (1:100), BV510 anti-IgD (1:100), FITC anti-IgM (1:100), PE anti-LAIR1 (1:100). After washing cells, SWM B cells defined as CD19^+^, CD27^+^, IgD^-^, IgM^-^ were isolated as LAIR1+ or LAIR1- by cell sorting and stored in TRIzol reagent (Life Technologies) at −80℃. RNA isolation, confirmation of RNA integrity, library preparation, and sequencing analysis were conducted at Azenta (AZENTA life sciences). Briefly, mRNA was selected with polyA selection from purified total RNA. 2×150 base pair, single index sequencing (paired-end) was performed using an Illumina HiSeq sequencer. Sequence reads were trimmed to remove possible adapter sequences and nucleotides with poor quality using Trimmomatic v.0.36. The trimmed reads were mapped to the Homo sapiens GRCh38 with ERCC genes reference genome available on ENSEMBL using the STAR aligner v.2.5.2b. Gene hit counts were calculated by using featureCounts (v.1.5.2). Using DESeq2, a comparison of gene expression between LAIR1+ and LAIR1- SWM B cells was performed. The Wald test was used to generate p-values and log2 fold changes. Genes with an adjusted p-value < 0.05 and absolute log2 fold change > 0.5 were called as differentially expressed genes for each comparison.

### Membrane Array for detection of phosphorylated LAIR1

B cells (2.0×10^6^ cells) isolated with EasySep human B cell isolation kit (Stem cell technologies) were rested in X-vivo medium (Lonza) at 37℃ for 30 min and subsequently stimulated with biotinylated anti-LAIR1 polyclonal antibody (1.0 μg/mL, R&D Systems) or biotinylated isotype antibody (1.0 μg/mL, R&D Systems) in the presence of streptavidin (10 μg/mL, R&D Systems) for 20 min. The reaction was stopped by adding ice cold HBSS and whole cell lysates were extracted with RIPA lysis buffer supplemented with Halt protease inhibitor cocktail (Thermo Fisher) and Halt phosphatase inhibitor cocktail (Thermo Fisher) for 30 min on ice. The lysates were incubated with human phosphor-immunoreceptor array membranes (R&D Systems) according to the manufacturer’s instructions. Phosphorylated proteins were detected by pan-anti-phospho-tyrosine antibody conjugated to HRP. Data were obtained using the Sapphire Biomolecular Imager (Azure Biosystems). Densitometric analysis of spot intensities was performed using Image J software. Phosphorylation levels of individual analytes were determined by averaging pixel densities of duplicate spots; values were normalized to positive controls, spot #1. Raw images of membrane arrays are shown in Supplementary Figure. 2.

### Phos-flow assay for pERK and pAKT

To avoid BCR activation with staining antibodies, naïve and unswitched memory (USWM) IgM+ B cells were defined as IgG- and IgA-. IgG+ B cells (SWM) were defined as IgM- and IgA-. Isolated B cells (2.0×10^6^ cells) were incubated with cocktails of Fixable Viability Dye eFluor 780 (1:1000), BV605 Anti-IgG (1:100) and PE-vio770 anti-IgA (1:100) for IgM+ B cells, or cocktails of Fixable Viability Dye eFluor 780 (1:1000), BV605 Anti-IgM (1:100) and PE-vio770 anti-IgA (1:100) for IgG+ B cells. After a washing step, the sorted cells were rested in X-vivo medium at 37℃ for 30 min and stimulated with biotinylated anti-LAIR1 polyclonal antibody (1.0 μg/mL, R&D Systems) or biotinylated isotype antibody (1.0 μg/mL, R&D Systems) in the absence or presence of streptavidin (5 μg/mL, Sigma-Aldrich) for 20 min. Cells were activated with biotinylated anti-IgM F(ab’)2 (3 μg/mL; Jackson ImmunoResearch), anti-IgM F(ab’)2 (3 μg/mL; Jackson ImmunoResearch), biotinylated anti-IgG F(ab’)2 (30 ng/mL; SouthernBiotech) or anti-IgG F(ab’)2 (30 ng/mL; SouthernBiotech) in the absence or presence of streptavidin (5 μg/mL). Cells were incubated for 10 min, and then further incubated with 5-fold volume (500 μL) of BD Cytofix™ Fixation Buffer (BD) for 10 min at 37°C. Cells were washed with FACS Buffer, and permeabilized in 200 µL of ice-cold Perm Buffer II (BD). After a washing step the cells were stained with anti-CD27, anti-LAIR1, anti-CD19, and anti-pAKT, and anti-pERK antibodies.

### Co-culture with cTfh cells and naïve B cells

Human CD4+ T cells were purified negatively from PBMCs using the EasySep CD4+ T cell isolation kit (Stem cell technologies) according to the manufacturer’s instructions. FVD-CD4+CD45RA-CXCR5+PD-1+ cells were sorted from these cells as circulating Tfh cells (cTfh cells).

Autologous naïve B cells were simultaneously isolated as FVD-CD3-CD14-CD16-CD56-CD123-CD27-IgA-IgG-cells from same donor.

Tfh cells (cTfh cells: 0.5×10^5^ cells) were co-cultured with autologous naïve B cells (1.0×10^5^ cells) at a ratio of 1:2 in 200 μL of X-vivo medium and stimulated with Dynabeads Human T-Activator CD3/CD28 Beads (0.5×10^6^ Beads; Gibco). Either recombinant human IL-21R Fc chimera protein (20 μg/mL; R&D Systems) or isotype control human IgG1 antibody (20 μg/mL; BioLegend) was added to cultures. After 5 days, cells were harvested and analyzed by flow cytometry. LAIR1 expression on B cells (CD3^-^, CD19^+^) were evaluated as geometric mean fluorescence intensity (Geo MFI) or the percentage of positive cells in B cell fraction.

### In vitro culture of naïve B cells

Sorted naïve B cells (2.0×10^5^ cells) were cultured in X-vivo medium for 3 days with various stimulations; anti-IgM F(ab’)2 (10 μg/mL; Jackson ImmunoResearch), MEGA CD40L (0.3 μg/mL; Enzo), CpG-B (1 μM; Invivogen), R848 (2.5 μg/mL; Invivogen), IL-21 (25 or 250 ng/mL; R&D Systems) and IL-10 (250 ng/mL; R&D Systems). In some experiments, 1 μM of SD36 was added to block STAT3. For differentiation to age associated B cells (ABCs), the differentiation cocktail, including anti-IgM F(ab’)2 (3 μg/mL), BAFF (10 ng/mL; Pepro tech), IL-2 (50 ng/mL; Pepro tech) and IFNγ (20 ng/mL; R&D Systems), was added to the culture medium with R848 or CpG-B and IL-21. Harvested cells were stained and analyzed by flow cytometry as described. T-bet was stained with Foxp3/Transcription Factor Staining Buffer Set (eBioscience). LAIR1 expression on CD19+ B cells was evaluated as Geo MFI or the percentage of positive cells in the total B cell fraction

### *In vitro* SWM B cell culture and plasmablast differentiation

LAIR1+ and LAIR1- CD19+CD27+IgD-IgM- SWM B cell (0.4×10^5^ cells) were cultured in X-vivo medium for 3 to 5 days under various stimulations; MEGA CD40L (0.3 μg/mL), CpG-B (1 μM), R848 (2.5 μg/mL), IL-21 (25 ng/mL) and IL-10 (250 ng/mL). Harvested cells were stained and analyzed by flow cytometry. PCs and Non-PCs were defined as CD27^++^, CD38^++^ and CD27^+^, CD38^mid^, respectively.

### Quantitative real-time PCR

Naïve B cells (0.6-2.0×10^6^ cells) were suspended in X-vivo medium at 37℃ for 30 min. They were stimulated with IL-21 (25 ng/mL) for 0, 2, 6 and 24 hours or IL-21 plus CpG-B or CD40L/Anti-IgM F(ab’)2 for 2 days. Cells were washed twice with ice cold HBSS and lysed with Qiazol (Qiagen). SD36 were treated prior to IL-21 stimulation for 3.5 hours. Lysates were freezed on −80℃ until the procedure of RNA isolation.

RNA was extracted from cells using RNeasy Mini Kit (Qiagen) according to the manufacturer’s instructions. Freshly isolated mRNA was reverse transcribed into complementary DNA with random primers using iScript cDNA Synthesis Kit (Bio-Rad).

Quantitative RT-PCR was performed on LightCycler 480 II (Roche) using TaqMan primers (Thermo Fisher Scientific), listed in Supplementary Table 3. Relative gene expression was normalized to a control gene, EIF1B eukaryotic translation initiation factor 1B (EIF1b), using the 2^-ΔΔCt^ method.

### Western blotting

Naïve B cells (6.0×10^6^ cells) were suspended in X-vivo medium at 37℃ water bath for 30 min. They were stimulated with IL-21 (25 ng/mL) for 30 min or with SD36 (1 μM) for 3.5 hours. Cells were washed twice with ice cold HBSS and whole cell lysates were extracted with RIPA buffer including the cocktail of protease inhibitors (Thermo) and phosphatase inhibitors (Thermo). Nuclear extracts were collected with NE-PER™ Nuclear and Cytoplasmic Extraction Reagents (Thermo) according to the manufacturer’s instructions. Whole lysates and nuclear extracts were obtained after centrifugation, and protein concentration was measured using Pierce™ Rapid Gold BCA Protein Assay Kit (Thermo). The lysates were analyzed by 4%–12% Bis-Tris PAGE and transferred to a PVDF membrane using iBlot2 (Invtrogen). The membrane was blocked for 1 hour at room temperature with 1% non-fat dry milk (Lab Scientific) in 0.1% Tween-20/Tris-buffered saline buffer (TBS-T) and then incubated with the primary antibodies overnight at 4°C. All primary antibodies were diluted (1:4000) with 5% BSA in TBS-T; anti-STAT1 (CST), anti-pSTAT1, pY701 (CST), anti-STAT3 (CST), anti-pSTAT3, pY705 (CST), anti-STAT5 (CST), anti-pSTAT5, pY694 (CST), anti-Lamin B1 (Abcam) and β-actin (CST). After three washes in TBS-T, the membranes were incubated for 1 h with anti-rabbit IgG, HRP-linked antibody in 5% BSA in TBS-T at 1:2000 (CST). After three washes in TBS-T, immunoreactive proteins were visualized with SuperSignal™ West Pico PLUS Chemiluminescent Substrate (Thermo). The membrane was scanned on a Sapphire Biomolecular Imager (Azure Biosystems). Full images of membranes are shown in Supplementary Figure. 6B, 6C and 6D.

### EMSA (Electrophoretic Mobility Shift Assay)

IRDye800-conjugated oligonucleotide was designed according to the LAIR1 promoter region sequence. The sequence used is shown in Supplementary Table 4.

Isolated naïve B cells using EasySep™ Human Naïve B Cell Isolation Kit (Stem Cell Technologies) were rested in X-VIVO medium for 30 min at 37℃ in 5% CO2 and subsequently stimulated with 25 ng/mL of human IL-21 (R&D Systems, #8879-IL-010/CF) for 30 min. Nuclear proteins were extracted with NE-PER™ Nuclear and Cytoplasmic Extraction Reagents (Thermo Fisher Scientific) according to the manufacturer’s instructions and stored at −80℃.

Upon EMSA, 3.0 μg of the defrosted nuclear extract were incubated with 7.5 μg of polyclonal anti-human STAT3 antibody (Sigma-Aldrich) or isotype control antibody for 20 min at room temperature and then incubated with the oligonucleotide for another 20 min using Odyssey EMSA Kit (LICOR bio) according to the manufacturer’s instructions. The oligonucleotide-nuclear extract complex was loaded to pre-run 5% TBE gel (Bio-Rad) and run at 100V for 50 min. The bands were visualized using Sapphire Biomolecular Imager (Azure Biosystems) and analyzed with ImageJ. Full images of membranes are shown in Supplementary Figure. 7.

### Terminology and B cell definition

Terminology and B cell definition in this study are summarized in Supplementary Table 6.

### Statistical analysis

Statistical analysis was performed using GraphPad Prism version 9 (Graph Pad) and IBM SPSS Statistics Version 26. *P* values <0.05 were considered statistically significant. The statistical method used in each experiment is reported in the figure legends.

### Study approval

The study with SLE patients was approved by the Northwell Health Institutional Review Board, and all subjects gave written informed consent.

## Results

### LAIR1 expression changes during activation and differentiation

We first characterized LAIR1 expression on human B cell subsets. Consistent with previous reports (14), LAIR1 expression on total B cells showed a clear biphasic pattern: The CD27^-^, IgD^+^ naïve B cell fraction was mainly LAIR1^+^ while CD27^+^ B cells and plasmablasts (PBs) contained the majority of LAIR1^-^ cells (Fig. 1A). Detailed analysis revealed that more than 99% of transitional B cells (TrB) and naïve B cells expressed LAIR1. LAIR1 positivity was approximately 83% for IgM^+^ CD27^+^ unswitched memory B cells (USWM), 60% for SWM B cells and 27% for PBs (Fig. 1B, Gating strategies are shown in Supplementary Figure. 1A). Furthermore, analysis of the expression levels on LAIR1^+^ cells showed that CD27^+^ B cells and PBs subsets had lower LAIR1 expression than TrB and naïve B cells (Fig.1B). Among SWM cells, IgA^+^ cells had a modest but significant decrease in LAIR1 positivity and expression level compared to IgG^+^ cells (Fig. 1C, the gating strategy is shown in Supplementary Figure. 1B). To further investigate the relationship between LAIR1 expression and B cell differentiation, we separated naïve B cells by CD45RB status. CD45RB denotes glycosylated epitope (17); glycosylated CD45RB^+^ (CD45RB^+^) naïve B cells are considered to be early memory B cells which are included among IgM^+^ USWM cells (18–20). Although these cells are a small population, we found that both LAIR1 positivity and the LAIR1 expression level on CD45RB^+^ naïve B cells were lower than on CD45RB^-^ naïve B cells (Fig. 1D, the gating strategy is shown in Supplementary Figure. 1C). Interestingly, CD45RB^+^ naïve B cells had two clear peaks for LAIR1 expression resembling the profile of USWM B cells and suggesting that CD45RB^+^ naïve B cells have an intermediate phenotype between naïve and memory B cells as previously suggested (20). Recently Glass et al. have identified CD95^+^ memory B cells as an activated subset of memory cells (20). We found that CD95^+^ SWM B cells had significantly less LAIR1 expression than CD95^-^ SWM B cells (Fig. 1D). LAIR1 positivity inversely correlated with CD95 expression. Approximately 84% of CD95^-^ SWM B cells were LAIR1^+^ compared to 34% of CD95^+^ SWM B cells. The frequency of LAIR1 positivity of CD95^+^ SWM B cells was similar to that of PBs (34%). These data in composite demonstrate that LAIR1 expression changes during B cell differentiation.

**Figure 1.**
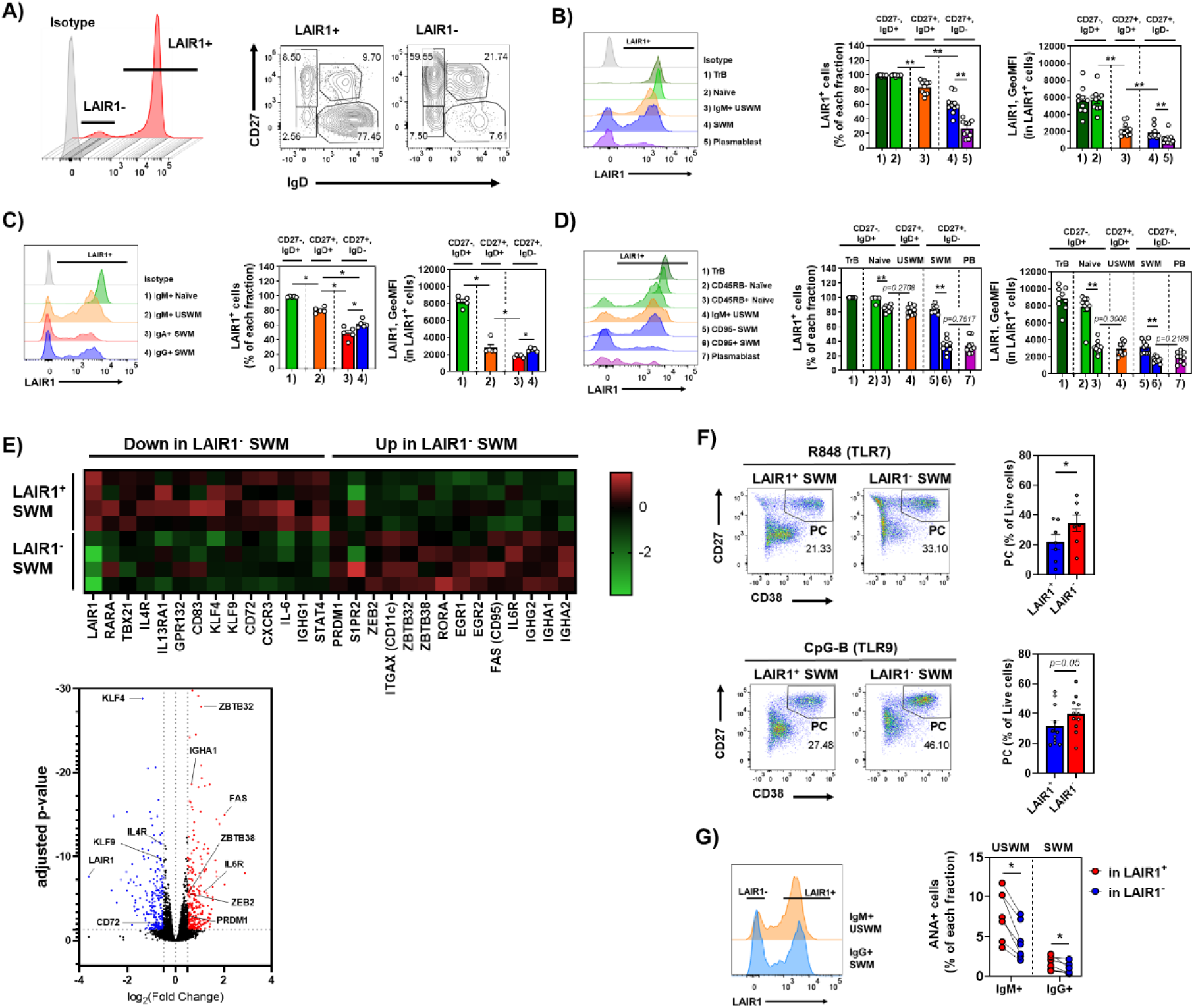
LAIR1 expression in various B cell subsets. (A) LAIR1 expression in total B cells (left) and the compositions of subsets in LAIR+ and LAIR1-B cells (right) (N=10). (B-D) LAIR1 expression in major B cell subsets (B; N=10), the different class of surface Ig (C; N=5) and major B cell subsets (D; N=9). (E) heatmap and volcano plot generated from RNA-seq data between LAIR1+ and LAIR1- SWM B cells (N=4). Differential gene expression was identified with Log_2_FC >0.5 and adjusted p value <0.05. Upregulated genes in LAIR1- are 278 genes and down-regulated genes in LAIR1- are 218 genes. (F) the potential of PC differentiation induced by TLR7 or TLR9 signaling. (G) the percentages of ANA+ autoreactive B cells in LAIR1+ and LAIR1- memory fraction (N=7). Data are shown as mean ± SEM with each symbol representing an individual subjects. P values were calculated with the Wilcoxon signed rank tests (F and G) and further adjusted for multiple comparisons using the Benjamini-Hochberg method (B, C and D). Asterisks indicate significant differences (\**P* < 0.05, \*\**P* < 0.01).

As LAIR1^-^ cells were more abundant in CD95^+^ SWM B cells, we hypothesized that LAIR1^-^ B cells may have a more activated phenotype. To address this hypothesis, we performed bulk RNA-seq on LAIR1^+^ and LAIR1^-^ SWM B cells. Transcriptomic analysis revealed 278 up-regulated genes and 218 down-regulated genes in LAIR1^-^ SWM B cells, some of which were related to B cell development and PC differentiation (Fig. 1E). For example, LAIR1+ SWM had B cell markers, *IL4R* (*21–23*), *KLF4* (*24*), *KLF9* (*24*) and *CD72* (*25, 26*), which are highly expressed in naïve B cells, while LAIR1^-^ SWM cells had more activated or differentiated markers, including *ZEB2* (*27*), *CD95* (*20, 25, 28*), *PRDM1* (*29*) and *IL6R* (*22*).

Because LAIR1^-^ SWM B cells had higher expression than LAIR1^+^ SWM B cells of *PRDM1* mRNA, a master transcription factor for PC differentiation (29–32) (Fig. 1E), we further investigated the differentiation potential of LAIR1^+^ and LAIR1^-^ SWM cells. As expected by the RNA-seq analysis, significantly more LAIR1^-^ SWM cells differentiated to PCs when stimulated with toll-like receptor (TLR) 7/8 ligand, R848, or TLR9 ligand, CpG-B, than LAIR1^+^ SWM cells (Fig. 1F). These data indicate that LAIR1^-^ SWM B cells have a more differentiated status than LAIR1^+^ SWM cells.

Given these data, we next examined LAIR1 expression on autoreactive B cells. Previously, we have shown that the frequency of anti-nuclear antigen reactive (ANA+) cells was higher in naïve B cells and declined as B cells matured to memory B cells (3, 4). We examined the frequency of ANA+ cells in LAIR1^+^ and LAIR1^-^ memory B cells. The LAIR1^+^ population contained more ANA+ cells compared to the LAIR1- population in both USWM and SWM B cells (Fig. 1G, the gating strategy of ANA is shown in Supplementary Figure. 1D). This suggests that LAIR1 may be a checkpoint on the activation of ANA+ memory cells in healthy individuals.

### LAIR1 can suppress BCR signaling

Some reports have suggested that LAIR1 suppresses BCR-induced Ca^2+^ influx and kinase cascades (14–16). We therefore evaluated the inhibitory function of LAIR1 on BCR signaling. We first confirmed that the anti-LAIR1 agonistic antibody was able to induce LAIR1 phosphorylation (pLAIR1) efficiently (Fig. 2A). Naïve, USWM B cells and SWM B cells were activated with biotinylated anti-IgM F(ab’)2 or biotinylated anti-IgG F(ab’)2 in the presence or absence of biotinylated anti-LAIR1 antibody and then exposed to streptavidin (Fig. 2B-D). LAIR1 engagement inhibited BCR-mediated ERK and AKT phosphorylation in all subsets only when BCR and LAIR1 were crosslinked with streptavidin, showing that LAIR1 limits BCR signaling when brought into proximity to the BCR.

**Figure 2.**
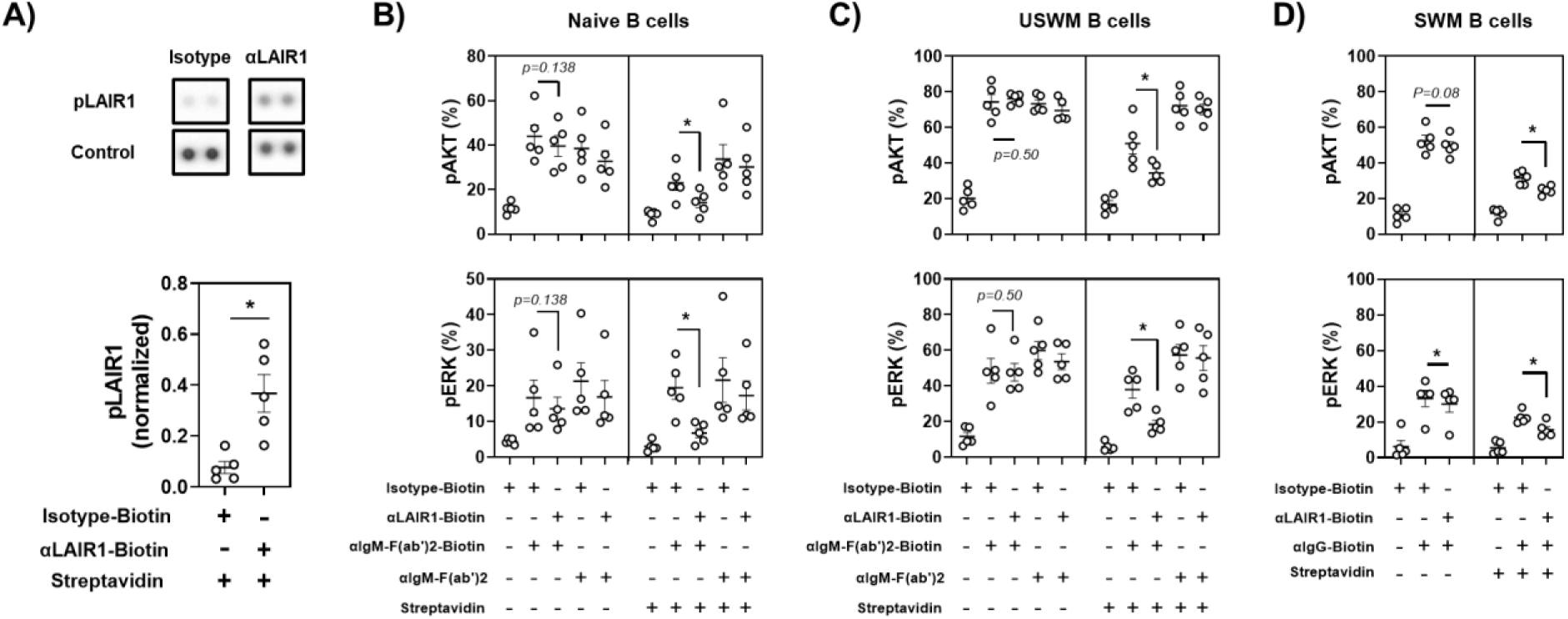
LAIR1 suppresses BCR signaling. (A) Phosphorylated LAIR1 induced by its crosslinking with biotinylated anti-LAIR1 antibody and streptavidin (N=5). (B-D) The effects of co-crosslinking with BCR and LAIR1 in naïve B cells (B; N=5), IgM+ USWM B cells (C; N=5) and IgG+ SWM B cells (D; N=5). Data are shown as mean ± SEM with each symbol representing an individual subjects. P values were calculated with the Wilcoxon signed rank tests. Asterisks indicate significant differences (\**P* < 0.05).

### Altered LAIR1 expression in SLE

The biased distribution of autoreactive B cells in the LAIR1^+^ compared to the LAIR1^-^ population led us to examine B cells from SLE patients. As shown in Fig. 3A, B cells of SLE patients shared the expression trends that we observed in age and sex matched healthy donors (Fig. 1); both the frequency of LAIR1^+^ cells and level of expression of LAIR1 decreased with differentiation (Fig. 3A, the gating strategy is shown in Supplementary Figure. 3A). However, we found reductions of LAIR1 positivity and expression level in several B cell subsets in the patients with SLE compared to healthy donors. Because of previous reports demonstrating differences in B cell subsets in SLE (33), we divided naïve B cells and CD27^-^ IgD^-^ double negative (DN) B cells into 4 subsets based on expression of CD21 and CXCR5: CD21^+^CXCR5^+^ resting naïve cells; CD21^-^CXCR5^-^ activated naïve cells; CD21^+^CXCR5^+^ DN (DN1) cells; and CD21^-^CXCR5^-^ DN (DN2) cells (The gating strategy is shown in Supplementary Figure. 3B, 3C). DN2 cells include T-bet^+^, CD11c^+^ Age associated B cell (ABC)-like cells which are reported to have a pathogenic role in SLE and transcriptome analysis and BCR repertoire studies suggest that activated naïve B cells are their precursor cells (33). Consistent with previous reports, we found an expansion of DN cells and decrease of USWM B cells in the patients, while frequencies of the other subsets were similar (Supplementary Figure. 3D and 3E). We observed LAIR1^-^ cells were expanded in the patients in all B cell populations except resting naïve B cells. Even though the frequencies of LAIR1 positivity in resting naïve B cells were comparable between healthy donors and SLE patients, LAIR1 expression level was significantly decreased (Fig. 3B). We then analyzed the frequency of LAIR1 positivity in B cells reactive with nuclear antigen (ANA+) in naïve, IgM^+^ USWM and IgG^+^ SWM B cells. ANA+ B cells had a modestly higher frequency of LAIR1^+^ cells compared to ANA-B cells in naïve and IgM^+^ USWM B cells both in healthy donors and SLE patients; interestingly, ANA+ B cells were not more frequently LAIR1^+^ in IgG^+^ SWM B cells of SLE patients (Fig. 3C). Furthermore, patients expressed a lower level of LAIR1 on ANA+ B cells than healthy donors (Fig. 3D).

**Figure 3.**
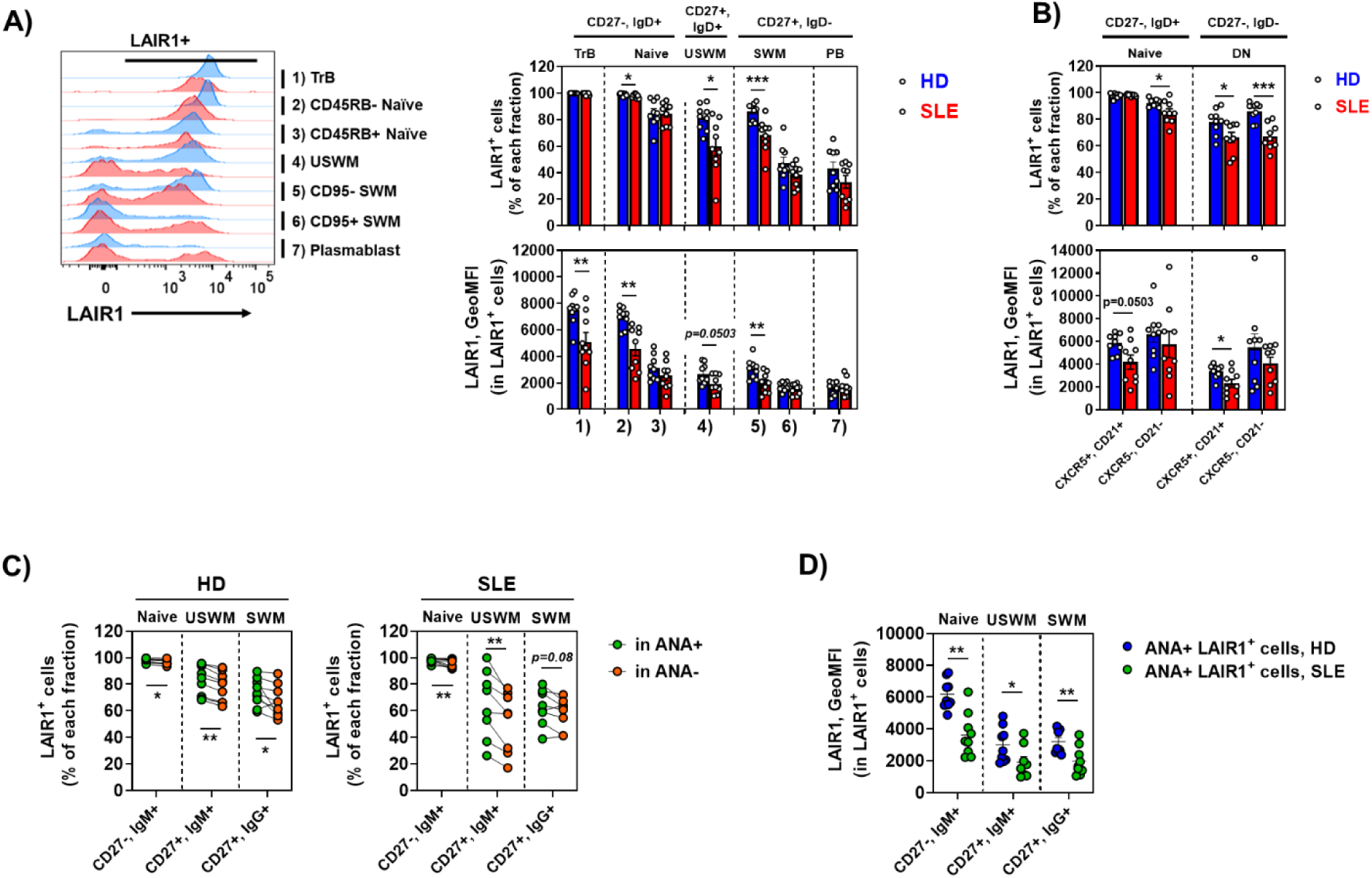
Impaired LAIR1 expression in SLE. (A-B) Representative examples of LAIR1 expression in 7 subsets of B cells (A) and activated naïve B cells and DN2 B cells (B) from HD and SLE patients (each N=9). (C) The positivity of LAIR1 in ANA+ and ANA- B cells. (D) The expression level of LAIR1 in LAIR1 expressing ANA+ autoreactive B cells between HD (N=9) and SLE patients (N=8). Data are shown as mean ± SEM with each symbol representing an individual subjects. P values were calculated with the Mann-Whitney U test (A, B and D) or the Wilcoxon signed rank tests (C). Asterisks indicate significant differences (\**P* < 0.05, \*\**P* < 0.01, \*\*\**P* < 0.001).

### IL-21 regulates LAIR1 expression

Given the reduced LAIR1 expression in patients with SLE, we sought to identify factors that downregulate its expression and hypothesized that they would be factors that are increased in SLE. The germinal center (GC) reaction is a crucial step during B cell activation and differentiation leading to the production of SWM B cells and PBs, these two subsets with reduced LAIR1 expression (Fig. 1B and 1D) and reported to contribute to autoantibody production in SLE (34, 35). In the light zone of GCs, B cells receive co-stimulatory signals such as IL-21 and CD40 ligand (CD40L) from T follicular helper (Tfh) cells as they differentiate into high-affinity memory cells and PBs (36–38). We, therefore, looked to see if IL-21 or CD40 engagement might downregulate LAIR1. First, we demonstrated that in vitro co-culture of naïve B cells with Tfh cells, activated by anti-CD3/CD28 coated beads, led to reduced LAIR1 expression (Fig. 4A). The reduction was diminished by the addition of an IL-21R Fc fusion protein which neutralizes IL-21 (Fig. 4A), suggesting that IL-21 is a regulator of LAIR1 expression. We confirmed the effect of IL-21 in cultures of naïve B cells stimulated with CD40L with or without BCR engagement (Fig. 4B). Naïve B cells stimulated with CD40L with or without BCR engagement increase expression of LAIR1 which is reduced in the presence of IL-21 (Fig 4B).

**Figure 4.**
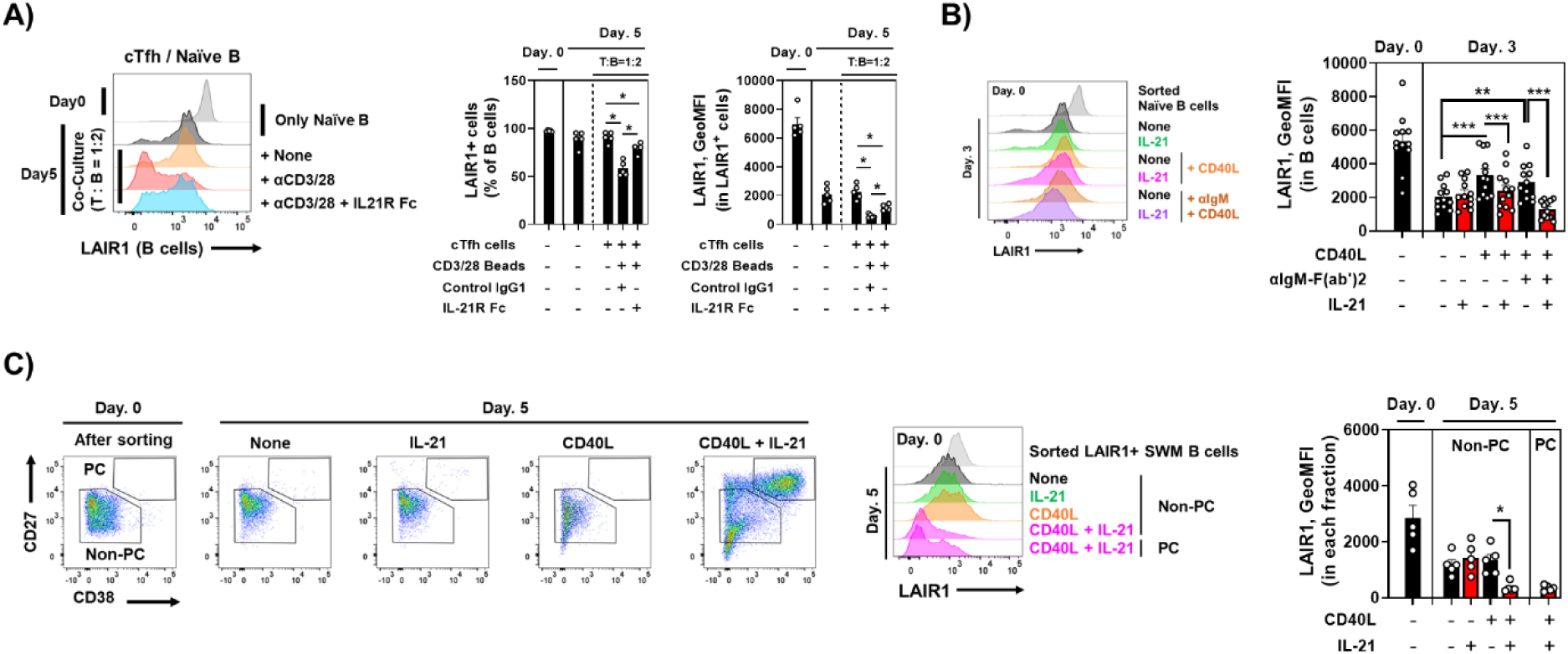
IL-21 decreases LAIR1 expression. (A) LAIR1 expression on B cells after 5 days co-culture with cTfh cells activated with anti-CD3/28 Beads in the presence of IL-21R-Fc protein (20 μg/mL) or isotype control (N=5). (B) The effects of IL-21 on LAIR1 expression in naïve B cells co-stimulated with CD40 or CD40/BCR in 3 days culture (N=12). (C) The effects of IL-21 on LAIR1 expression in SWM B cells co-stimulated with CD40 in 5 days culture (N=5). Data are shown as mean ± SEM with each symbol representing an individual subjects. P values were calculated with the Wilcoxon signed rank tests (C) and further adjusted for multiple comparisons using the Benjamini-Hochberg method (A and B). Asterisks indicate significant differences (\**P* < 0.05, \*\**P* < 0.01, \*\*\**P* < 0.001).

Activation of CD40 and IL-21 also induces PC differentiation from SWM B cells (36, 39, 40). When LAIR1^+^ SWM B cells were cultured with CD40L and IL-21, we observed a reduction in LAIR1 expression after 5 days of culture on both PCs and non-PCs (Fig. 4C). Thus, IL-21 regulates LAIR1 expression not only in naïve B cells, but also in SWM B cells.

### IL-21 regulates LAIR1 expression during extrafollicular B cell maturation

The extrafollicular (EF) pathway is another important mode of B cell activation and differentiation which has been implicated in SLE pathogenesis (2, 33). We, therefore, tested whether IL-21 also regulates LAIR1 expression in the presence of TLR agonists. In vitro stimulation of naïve B cells with CpG-B with or without BCR engagement increased LAIR1 expression; the addition of IL-21 at 25 ng/mL reduced LAIR1 expression (Fig. 5A, supplemental Fig. 5A). IL-21 at 25 ng/mL did not decrease LAIR1 expression in naïve B cells stimulated with R848; however, a significant reduction was observed in the presence of IL-21 at 250 ng/mL (supplemental Fig. 5B), suggesting stronger IL-21 signaling is required to reduce LAIR1 expression in TLR7 activated healthy naïve B cells.

**Figure 5.**
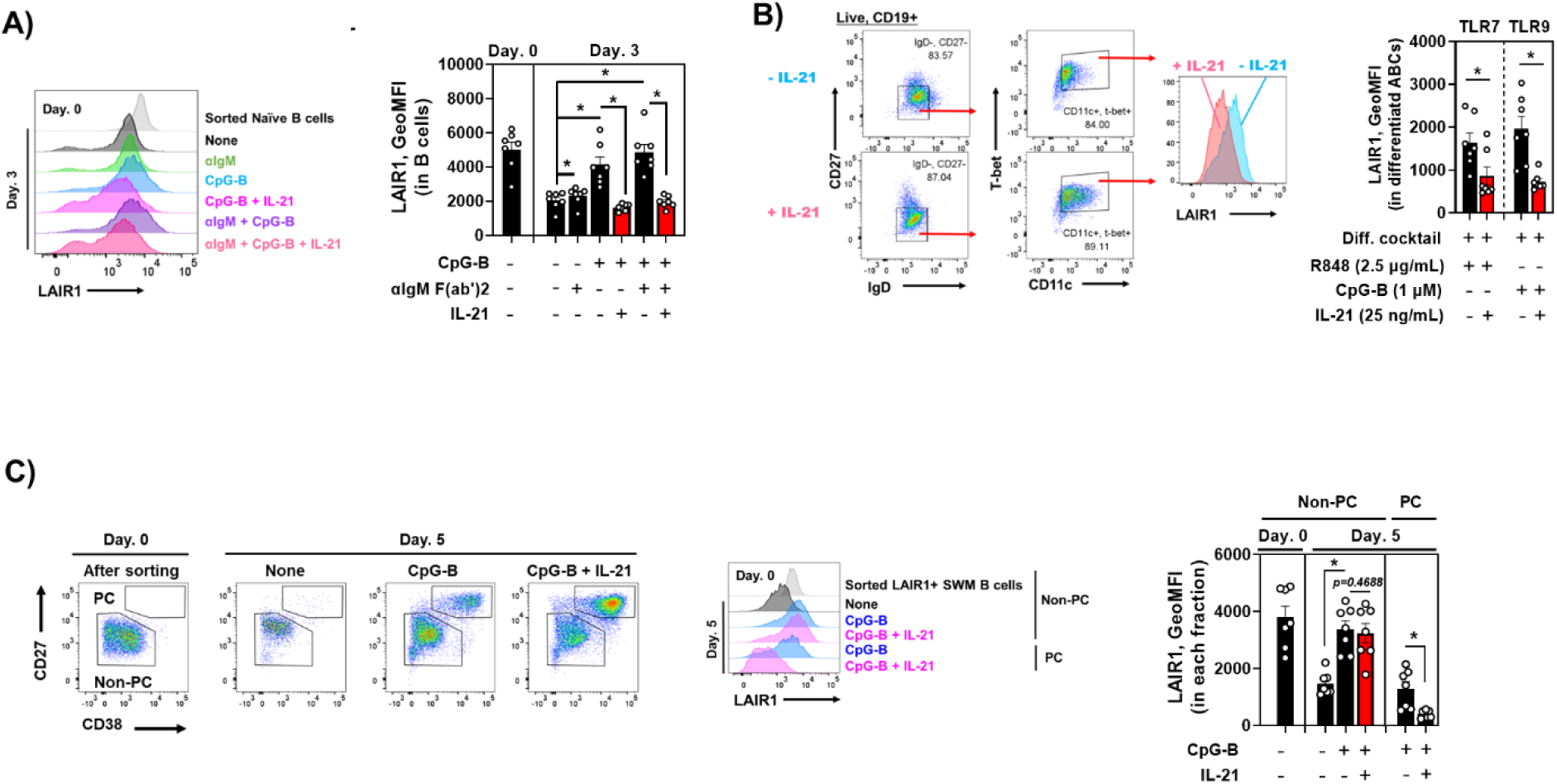
IL-21 decreases LAIR1 expression induced by TLR9. (A) The effects of IL-21 on LAIR1 expression in naïve B cells co-stimulated with TLR9 or TLR9/BCR in 3 days culture (N=7) (B) The effects of IL-21 on LAIR1 expression in ABCs differentiation from naïve B cells in 3 days culture (N=7). (C) The effects of IL-21 on LAIR1 expression in SWM B cells co-stimulated with TLR9 in 5 days culture (N=7). Data are shown as mean ± SEM with each symbol representing an individual subjects. P values were calculated with the Wilcoxon signed rank tests (B) and further adjusted for multiple comparisons using the Benjamini–Hochberg method (A and C). Asterisks indicate significant differences (\**P* < 0.05).

Ester et al. showed interferon γ (IFNγ) signaling induced epigenetic programming of B cells and accelerated TLR7 and IL-21 responses. IFNγ can also induce T-bet+, CD11c+ ABC-like cells from naïve B cells (33, 41). Human ABCs have a pathogenic role in SLE (33, 42) and are reported to undergo EF differentiation (33, 43). We, therefore, examined the expression of LAIR1 in naïve B cells induced to become ABC-like cells using a cocktail of IFNγ, IL-2, BAFF, anti-IgM F(ab’)2 and a TLR 7 or 9 agonist in the presence or absence of IL-21. LAIR1 expression was reduced on ABC-like cells induced with either CpG-B or R848 when IL-21 was present (Fig. 5B). We also tested LAIR1 expression on PCs differentiated from LAIR1^+^ SWM cells by CpG-B with or without IL-21 (Fig. 5C). Distinct to CD40L, CpG-B alone can induce PC differentiation from SWM B cells. PCs differentiated by CpG-B in the presence of IL-21 expressed less LAIR1 than those differentiated without IL-21. Similar results were observed when SWM B cells were stimulated with R848 in the presence of IL-21 (Supplemental Fig. 5C). Thus, IL-21 negatively regulates LAIR1 expression in B cells activated through an EF-like pathway.

### IL-21-induced STAT3 regulates LAIR1 promoter activity

Next, we examined whether IL-21 alters LAIR1 transcription in stimulated naïve B cells (Fig. 6A). As expected, IL-21 decreased LAIR1 mRNA expression in naïve B cells stimulated with TLR9 or CD40L plus BCR engagement. IL-21 decreased LAIR1 mRNA expression by itself. Reduction of LAIR1 mRNA was observed after 6 hours and sustained for at least 2 days (Fig. 6A and 6B), while its protein expression was not affected in this time frame, suggesting that another factor or more time is required for the decreased protein expression.

**Figure 6.**
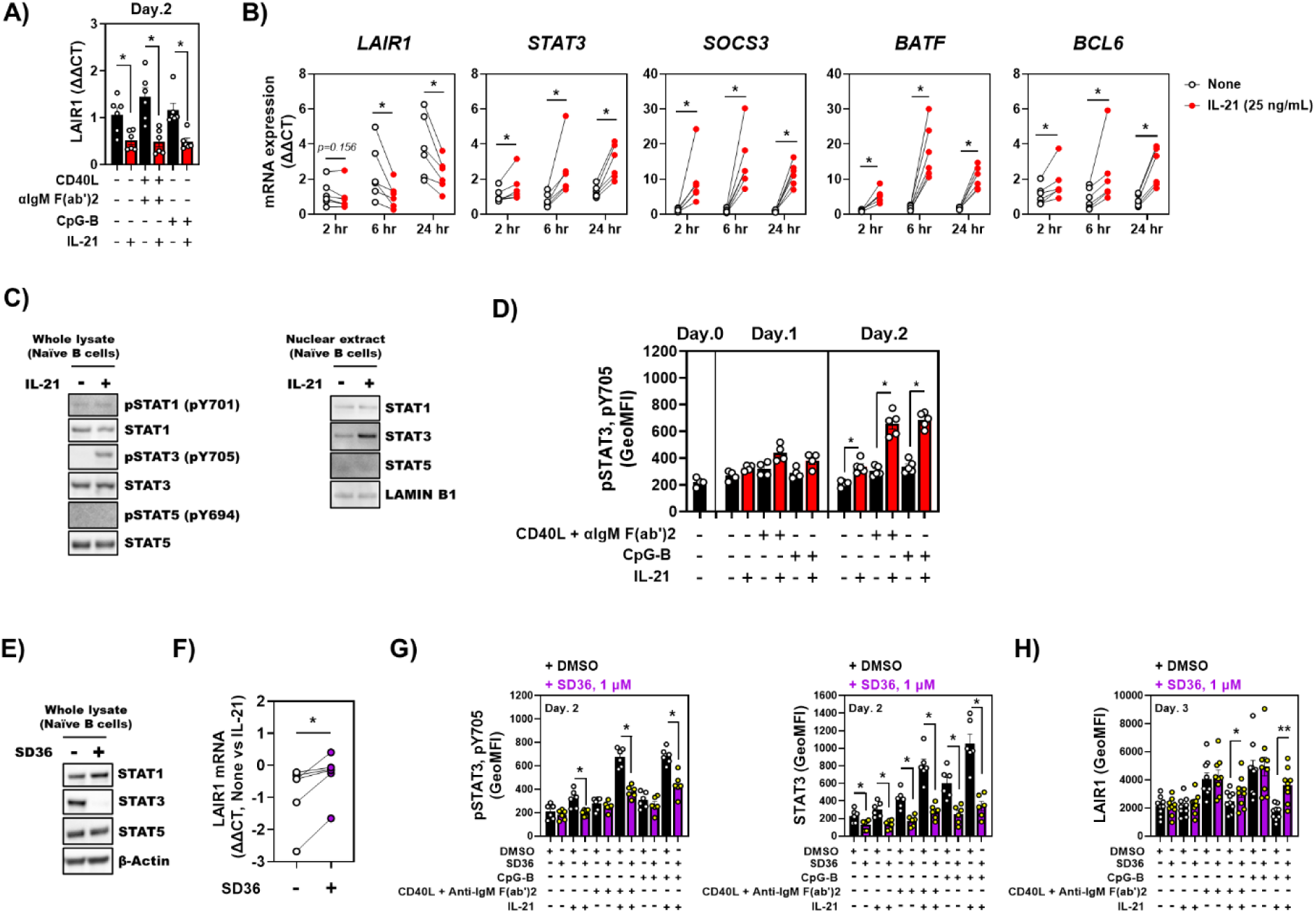
phospho-STAT3 is associated with decreased LAIR1 expression on naïve B cells. (A) The effects of IL-21 on LAIR1 mRNA expression in naïve B cells co-stimulated with TLR9 or CD40/BCR in 2 days culture (N=6) (B) The effects of IL-21 on LAIR1 mRNA expression in naïve B cells in 2-24 hours culture (N=7). (C) IL-21-induced phosphorylation (left) and nuclear translocation (right) of STAT1/3/5 (N=3). (D) Amplification of IL-21-induced pSTAT3 level by TLR9 or CD40/BCR in the naïve B cell culture (N=4-5). (E-G) the effect of SD36 (1 μM) against STAT family proteins for 3.5 hours (E; N=3), LAIR1 mRNA expression change induced by IL-21 (F; N=6), pSTAT3 and the protein levels of STAT3 (G; N=6) and LAIR1 (H; N=11). Data are shown as mean ± SEM with each symbol representing an individual subjects. P values were calculated with the Wilcoxon signed rank tests (A, B, D, F and G). Asterisks indicate significant differences (\**P* < 0.05, \*\**P* < 0.01).

IL-21 activates a JAK/STAT pathway and induces phosphorylation of STAT1, STAT3 and STAT5 (40, 44). We, therefore, investigated whether STAT family proteins regulated LAIR1 expression. First, the single stimulation of IL-21 induced not only STAT3 mRNA, but also transcriptional STAT3 target genes (Fig. 6B). Moreover, consistent with previous reports, IL-21 induced phosphorylation of STAT3 (pSTAT3) and translocation to the nucleus; both phosphorylation and nuclear translocation of STAT1 and STAT5 were not induced by IL-21 in our assay (Fig. 6C and D). To investigate whether IL-21-induced STAT3 negatively regulated LAIR1 expression, we treated naïve B cells with SD36, PROTAC compound for STAT3. SD36 is composed of 2 motifs: a STAT3-binding motif and a ubiquitin ligase (Celebron)-binding motif ligated with a linker which degrades STAT3 in a proteasome-dependent manner (45, 46). We confirmed by Western blot that SD36 clearly decreased STAT3 protein expression in 3.5 hours (Fig. 6E). IL-21 decreased LAIR1 mRNA expression and amplified the amount of STAT3 mRNA and phosphorylation of STAT3 in naïve B cells activated either by CD40 with BCR signaling or by TLR signaling (Fig. 6A, Fig. 6B, 6D and supplemental Fig. 6A), which is consistent with previous reports that STAT3 induces itself by binding its promoter region to create a positive feedback loop (47–50). Pretreatment of B cells with SD36 suppressed the IL-21 mediated decrease of LAIR1 mRNA as well as increase of STAT3-direct target genes, such as SOCS3 and BATF (Fig. 6F and supplemental Fig. 6B). We also confirmed that SD36 decreased both total and phosphorylated STAT3 (Fig. 6G), and rescued LAIR1 expression on IL-21 stimulated B cells (Fig. 6H), suggesting that STAT3 is a dominant regulator of LAIR-1 expression.

We next investigated whether STAT3 directly regulates LAIR1. Three potential STAT3 binding sites were identified within 1500 base pairs of the start site (Fig. 7A). Within these 3 sequences, only sequence #3 exists in LAIR1 proximal promoter region. We conducted an electrophoretic mobility shift assay (EMSA) for sequence #3 to examine if STAT3 directly binds to the sequence *in vitro*. As shown in Fig. 7B and C, a protein/DNA complex was detected using nuclear extract (NE) of IL-21 treated naïve B cells and the sequence #3 oligonucleotide and the addition of anti-STAT3 antibody diminished the band, implying that STAT3 in the NE bound to the sequence. Thus, STAT3 can bind the LAIR1 promoter region.

**Figure 7.**
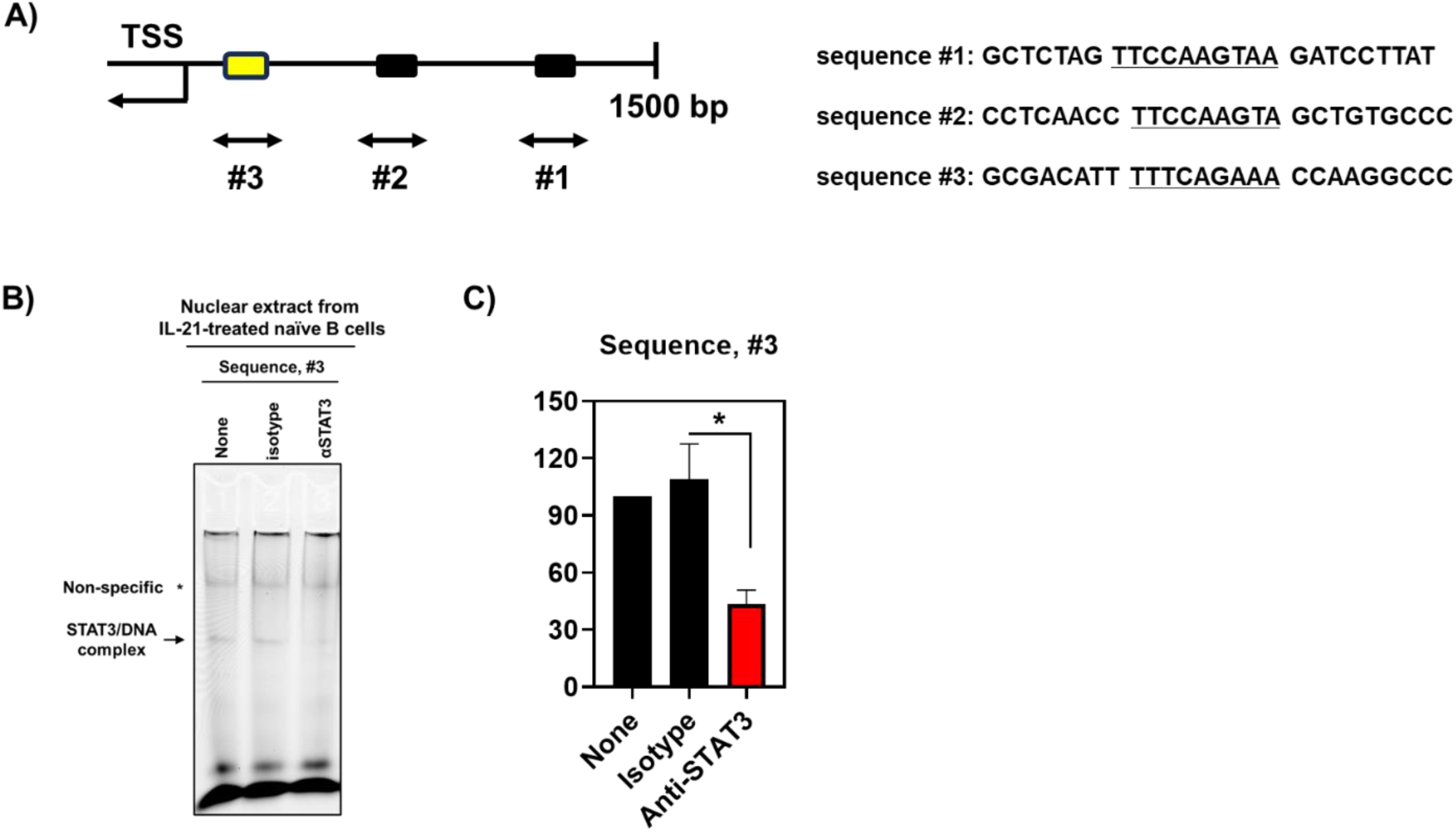
STAT3 directly binds the promoter region of LAIR1. (A) STAT3 putative binding site in LAIR1 promoter region and its sequence (B) the binding of STAT3 of nuclear extract from IL-21 treated naïve B cells to LAIR1 promoter region (sequence #3) (N=3). P values were calculated with the Paired T test (C). Asterisk indicates significant difference (\**P* < 0.05).

IL-10 is also known to activate STAT3. We, therefore, asked whether IL-10 might decrease LAIR1 expression. We cultured naïve B cells with CD40L and anti-BCR with and without IL-10. IL-10 decreased LAIR1 expression but to a lesser extent than IL-21 (supplemental Fig 4). There was no additive effect of IL-10 and IL-21, suggesting that they both operated through the same pathway.

## Discussion

In this study, we analyzed the expression profile of the inhibitory receptor, LAIR1, on human B cells. We demonstrated that LAIR1 expression gradually decreases during activation and differentiation, implying a relationship between LAIR1 expression and B cell development. In patients with SLE, the expression of LAIR1 was decreased and the frequency of LAIR1^-^ cells was expanded in several B cell subsets. The LAIR1^+^ fraction harbored more ANA+ autoreactive B cells than the LAIR1^-^ fraction, suggesting that LAIR1 is a tolerance checkpoint. We further identified a mechanism for downregulation of LAIR1 expression; the IL-21/STAT3 axis works as a negative regulator of LAIR1 in B cells by directly binding the LAIR1 proximal promoter region.

We have confirmed that co-localization of LAIR1 with BCR diminishes the strength of the BCR signaling. The known ligands for LAIR1 are collagen (9) and C1q (10). It is not obvious that collagen could be a means to bring LAIR1 and the BCR into proximity. C1q, however, can bind to immune complexes; antigen in the immune complexes could bind the BCR and C1q which might then bind LAIR1. Thus, just as IgG immune complexes colligate the BCR and FcRγ2b to suppress BCR signaling (51, 52), both IgM and IgG immune complexes might colligate the BCR and LAIR1.

PCs can be differentiated through either an EF pathway or as part of a GC response. Whether LAIR1 actually limits PC differentiation, or alternatively marks a B cell state that is less easily differentiated to PCs is not clear. In these studies, there was no clear ligand for LAIR1. Thus, there would need to be some tonic LAIR1 signaling for LAIR1 to be an actual checkpoint on PC differentiation. We believe the latter is more likely as the transcriptional analysis of LAIR1^+^ and LAIR1^-^ SWM B cells showed LAIR1^-^ B cells were more differentiated to a PC transcriptional profile.

Once naïve B cells recognize antigen and receive repetitive T cell help in GCs, LAIR1 expression is down-regulated potentially facilitating their differentiation to PCs. We attempted to stimulate a GC-like response by incubating B cells with Tfh cells or with CD40L and IL-21, both of which are critical in the GC response. We demonstrated that IL-21 increased PC differentiation in SWM B cells stimulated with CD40L. The GC response is a crucial step not only for affinity maturation, but also for the elimination of autoreactive B cells peripherally (53). We previously found that the frequency of ANA+ cells was high in naïve B cells and declined as B cells matured into memory B cells (3, 4). We demonstrated that the LAIR1^+^ memory B cell population contained more ANA+ cells than the LAIR1^-^ population (Fig. 1F), demonstrating a relationship between LAIR1 expression and ANA+ autoreactivity. Cappione et al. have shown that autoreactive B cells bearing the 9G4 idiotype are barely seen in GCs in healthy human tonsils (53). It is interesting to speculate that during the GC response, autoreactive B cells are eliminated and non-autoreactive and antigen-specific SWM B cells and PCs emerge with low to no LAIR1 expression. Thus, LAIR1^-^ SWM cells which have a more differentiated phenotype and more potential to become PCs than LAIR^+^ SWM cells may be generated through the GC response. In addition to the GC pathway, PCs can be differentiated through an EF pathway with endosomal TLR activation; this is thought to be an important pathway for PC differentiation in SLE (33, 43). In fact, Tipton et al. demonstrated that the repertoire of autoreactive PCs had clonal similarities to naïve cells rather than IgG^+^ memory B cells, suggesting that these PCs may rise through an EF pathway (43). Although it is still controversial which pathway is dominant for ANA+ PC differentiation in SLE, IL-21 can facilitate B cell activation and differentiation in both pathways (33, 42).

Several observations suggest the relevance of IL-21 in SLE, such as IL-21/IL-21 receptor risk alleles (54, 55), high serum/plasma levels of IL-21 (56, 57) and higher IL-21 production from CD4+ T cells of SLE patients (58, 59). We found that LAIR1 expression on B cells of SLE patients was significantly lower, including on ANA+ cells (Fig. 3) and IL-21 decreased LAIR1 expression upon co-stimulation with TLR7/9 in vitro (Fig. 4, Fig. 5). Thus, in B cells activated through endosomal TLRs, IL-21 may down-regulate LAIR1 expression and thereby facilitate the extrafollicular differentiation of ANA+ B cells to PCs. It is interesting to note that STAT3 deficient individuals have reduced USWM and SWM B cells supporting the relevance of STAT3 to both a GC and an EF pathway of B cell differentiation (40).

In 1999, Linde and her colleagues reported LAIR1 expression on human B cells for the first time (14). They showed a distinct expression profile among naïve B cells, GC B cells, memory B cells and PBs in tonsil and bone marrow PCs. Their data are largely consistent with our present study. LAIR1 was highly expressed in naïve B cells but was absent on approximately 50% of memory B cells and all PBs/PCs. The only difference between their data and the present study lies in LAIR1 positivity in PBs/PCs. While they showed that all PBs/PCs from tonsil and bone marrow have no expression of LAIR1, we found that 20% of PBs in blood expressed it. Blood PBs are composed of GC pathway-derived and EF pathway-derived cells. We speculate that EF pathway-derived PBs express more LAIR1 compared to the GC-derived PBs. In our *in vitro* experiments, TLR7- or TLR9-induced PCs maintained more LAIR1 expression than PCs induced by TLR or CD40 activation in the presence of IL-21. EF-derived PCs that differentiate without T cell help, in the absence of IL-21 may, therefore, maintain LAIR1 expression. Van der Vuurst de Vries et al. found no LAIR1 expression on GC B cells (CD38+, IgD-) and PBs (CD38++, IgD-) in human tonsils (14), suggesting that GC pathway-derived PCs may not express LAIR1 *in vivo*, which aligns to our data. Several studies have shown the clear differences of gene expression profiles between blood PBs and bone marrow PCs (BMPCs) (60–63), and single cell RNA-seq analysis for bone marrow PCs suggests that there are maturation steps for long lived PCs (LLPCs) in bone marrow (64). It is unclear whether this maturation process affects LAIR1 expression or whether LAIR1 is reduced prior to the final maturation phase.

Although previous reports demonstrated that TLRs and cytokines could affect LAIR1 expression in several cell types (65–67), no clear mechanism has been mentioned. We demonstrated that IL-21/STAT3 axis negatively regulates LAIR1 expression in human naïve B cells by binding to its proximal promoter. For human naïve B cells, Van der Vuurst de Vries et al. showed that BCR or CD40 stimulation can trigger a decrease in LAIR1 expression, and these effects were enhanced with IL-2 and IL-10 (14). We did not observe that CD40L or BCR signaling decreased LAIR1 expression. It is possible that different experimental conditions account for the discrepancy: They cultured naïve B cells from tonsils and analyzed LAIR1 expression on day 6, while we studied blood naïve B cells on day 3. In their cultures, they found LAIR1 reduction starting on day 5. IL-10 also activates STAT3 in B cells, presumably accounting for the IL-10 induced decrease in LAIR1. However, the effect of IL-10 was weaker than that of IL-21 in our studies (supplemental Figure. 4), suggesting that IL-21 is more dominant factor than IL-10 for the regulation of LAIR1 expression.

Unexpectedly, we found that IL-21 stimulation decreased LAIR1 mRNA at 6 hours, but not protein expression, while IL-21 together with TLR9 or CD40/BCR signaling decreased both LAIR1 mRNA and protein expression (Fig. 4B, 5A and 6D). Considering that IL-21 induced STAT3 binding on the proximal region of LAIR1 promoter, it suggests that another factor may be required for protein reduction. Protein levels in cells are affected by the rate of cell proliferation and the balance of protein synthesis and degradation. IL-21 is known to induce proliferation and a metabolically active phenotype in combination with TLR9 or CD40/BCR (39). Thus, it is plausible to think that IL-21 by itself did not decrease LAIR1 protein expression *in vitro* because of less cell proliferation and less protein turnover. In contrast, co-stimulation of IL-21 with TLR9 or CD40/BCR would enhance proliferation and decrease protein expression. It is also possible that TLR9 and signaling through CD40 and the BCR induces other factors that help to decrease the LAIR1 protein stability.

In conclusion, we found that IL-21/STAT3 axis is a dominant pathway for LAIR1 down-regulation during B cell differentiation. This axis may contribute to aberrant PC differentiation and the pathogenesis of SLE.

## Author Contribution

KY conceptualized the study, designed and conducted experiments, analyzed data, and wrote the manuscript. IKS conducted experiments, analyzed data and wrote the manuscript. YAF designed statistical analysis. CA conducted experiments. BD conceptualized the study, designed experiments, interpreted the data, and wrote the manuscript.

## Acknowledgements

The authors thank Guangchun Jin, Christopher Colon, Max Wallach and Michael Labarbera for technical assistance of flow cytometry and cell sorting. We also thank Myoungsun Son and Sun Jung Kim for helpful comments. The work was supported by a grant from the NIH (P01 AI148102 NIAID HHS United States) and funds to Kenji Yoshida from Asahi Kasei Pharma Corporation JAPAN)

