## Supplemental Figures 1-7 and Tables 1-6 for "IL-21-STAT3 axis negatively regulates LAIR1 expression in B cells"

**Supplemental Fig. 1**

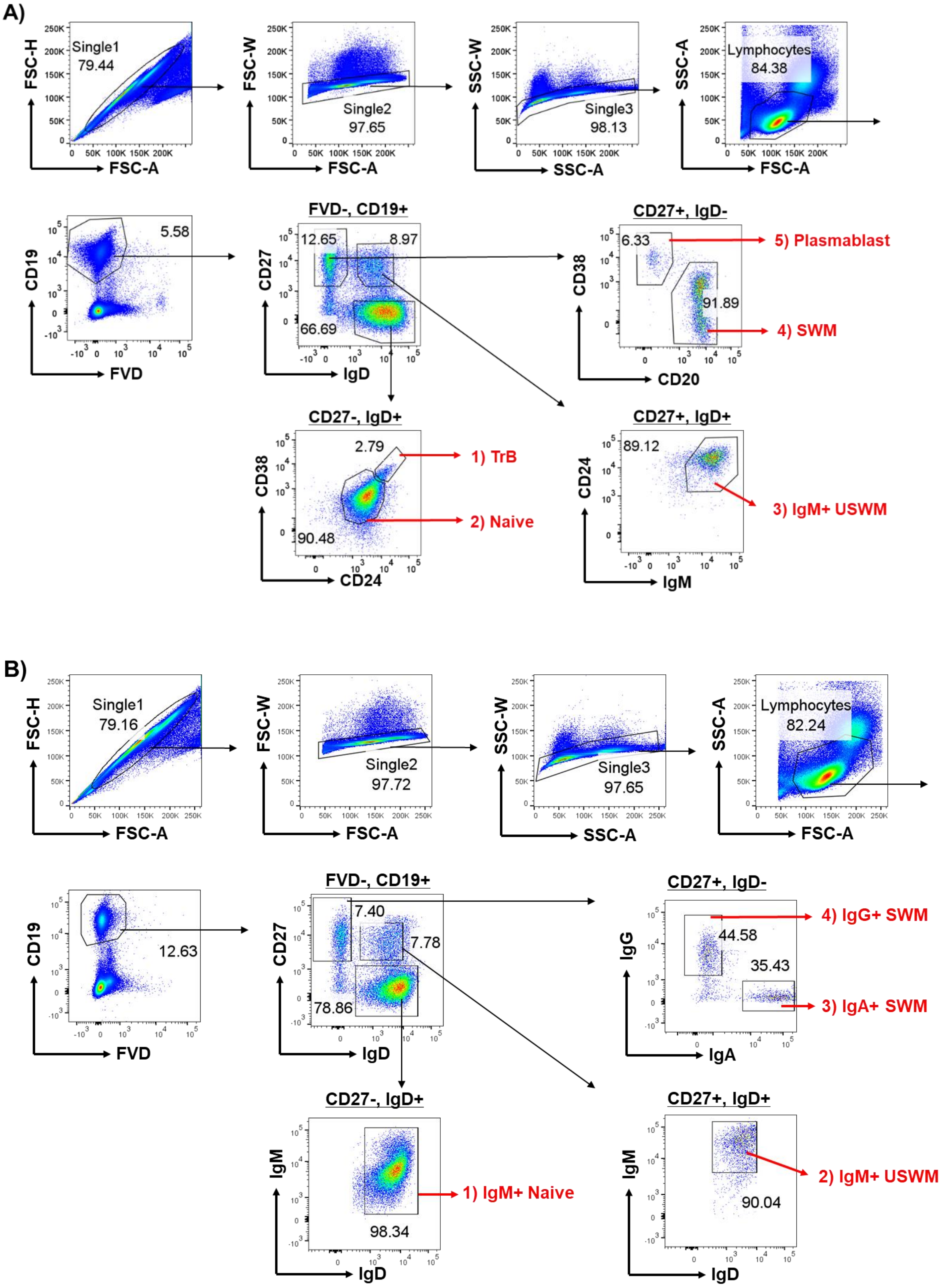

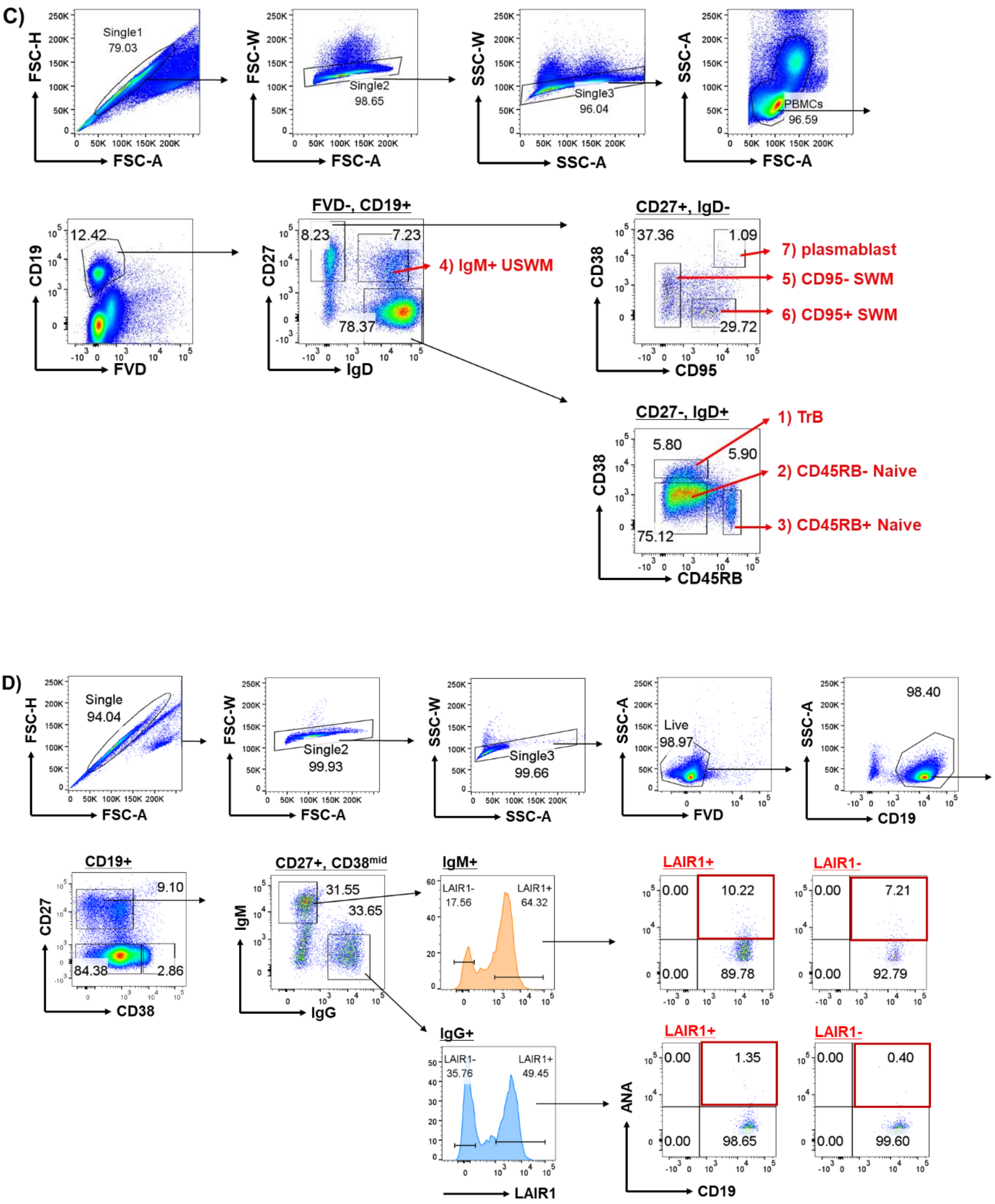


(A) The gating strategy for Figure 1B. (B) The gating strategy for Figure 1C. (C) The gating strategy for Figure 1D. (D) The gating strategy for Figure 1G.

### Supplemental Fig. 2

A)

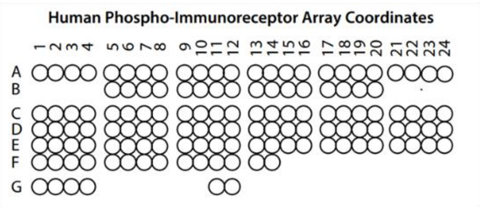


B)

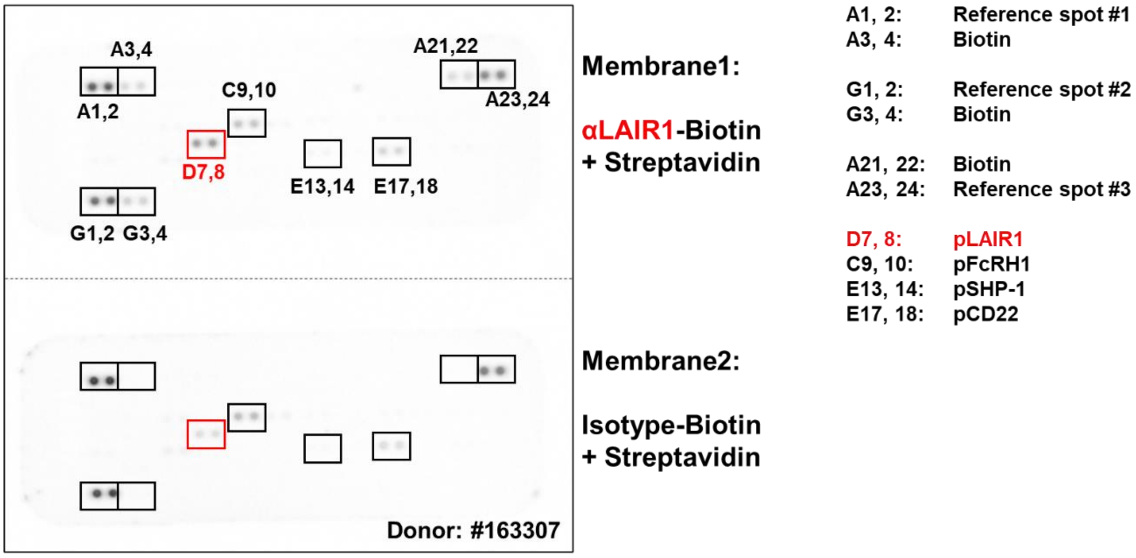


(A) Reference of Human Phospho-Immunoreceptor Array coordinates. (B) Full images of membrane arrays incubated with lysates from human B cells treated with biotinylated anti-LAIR antibody (Membrane1) or biotinylated isotype control antibody (Membrane2) in the presence of streptavidin.

### Supplemental Fig. 3

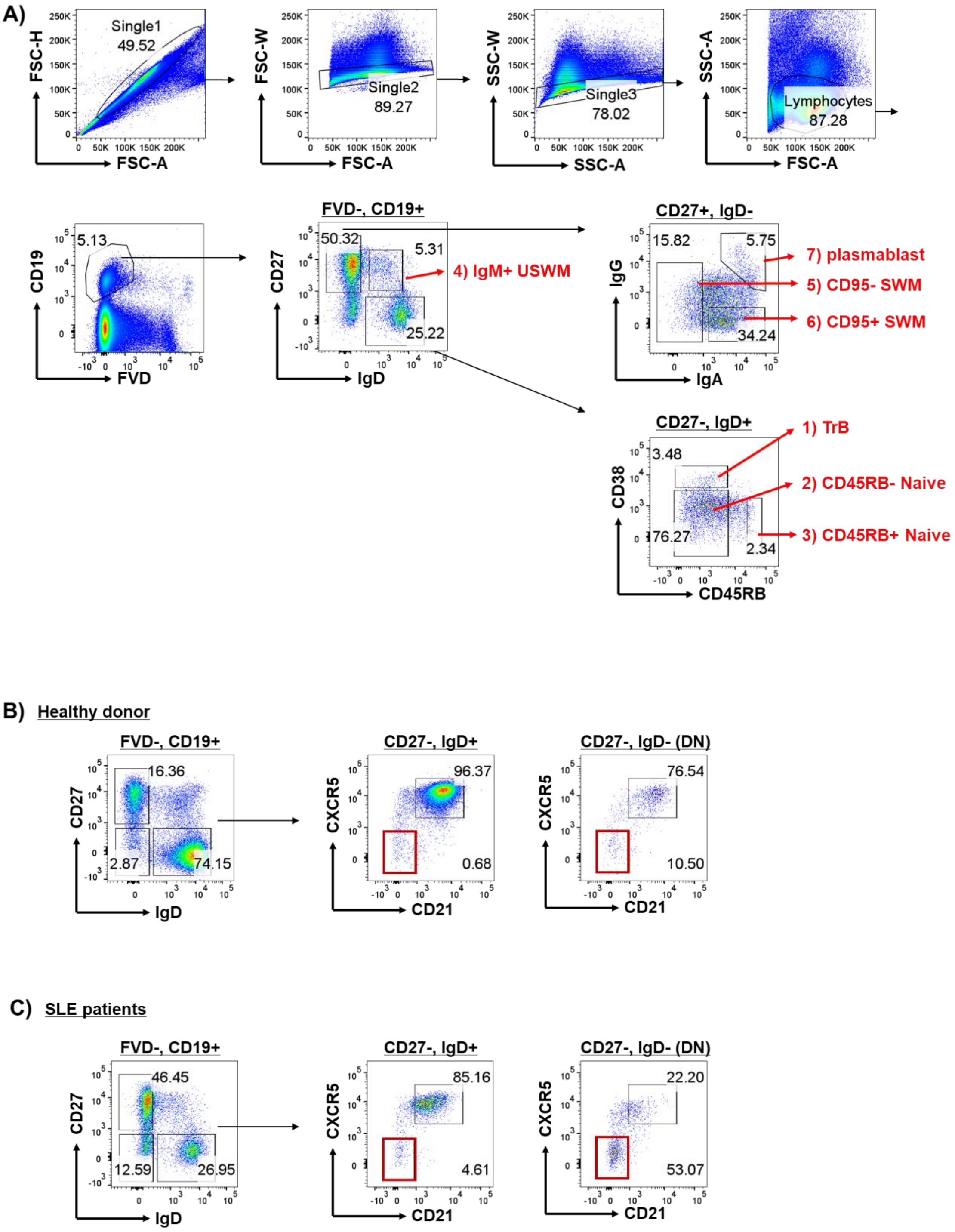

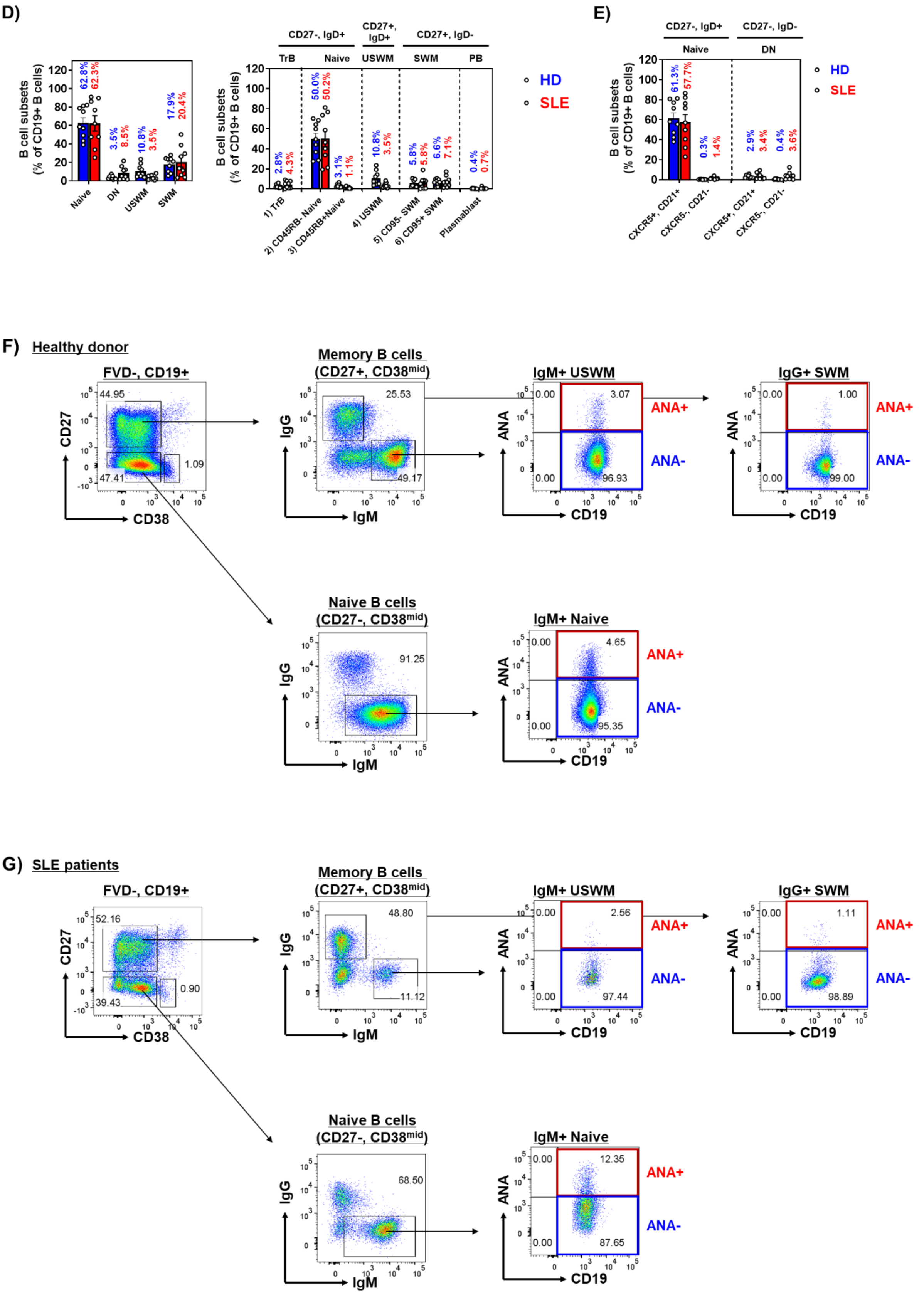

D)
B cell subsets (% of CD19+ B cells)
Naive
DN
USWM
SWM
62.8%
62.3%
3.5%
8.5%
10.8%
3.5%
17.9%
20.4%
CD27-, IgD+
CD27+, IgD+
CD27+, IgD-
TrB
Naive
USWM
SWM
PB
1) TrB
2) CD45RB- Naive
3) CD45RB+Naive
4) USWM
5) CD95- SWM
6) CD95+ SWM
Plasmablast
2.8%
4.3%
50.0%
50.2%
3.1%
1.1%
10.8%
3.5%
5.8%
5.8%
6.6%
7.1%
0.4%
0.7%
HD
SLE
E)
CD27-, IgD+
CD27-, IgD-
Naive
DN
CXCR5+, CD21+
CXCR5-, CD21-
61.3%
57.7%
0.3%
1.4%
2.9%
3.4%
0.4%
3.6%
F) Healthy donor
FVD-, CD19+
CD27
CD38
44.95
47.41
1.09
Memory B cells (CD27+, CD38mid)
IgG
IgM
25.53
49.17
IgM+ USWM
ANA
CD19
0.00
3.07
96.93
ANA+
ANA-
IgG+ SWM
1.00
99.00
Naive B cells (CD27-, CD38mid)
91.25
IgM+ Naive
4.65
95.35
G) SLE patients
FVD-, CD19+
52.16
39.43
0.90
Memory B cells (CD27+, CD38mid)
48.80
11.12
IgM+ USWM
2.56
97.44
IgG+ SWM
1.11
98.89
Naive B cells (CD27-, CD38mid)
68.50
IgM+ Naive
12.35
87.65

(A) The gating strategy for Figure 3A. (B, C) The gating strategy for Figure 3B (B: PBMCs from healthy donors, C: PBMCs from patients with SLE). (F, G) The frequencies of B cell subsets from PBMCs (D: Data from samples of Figure 3A, E: Data from samples of Figure 3B). (F, G) The gating strategy for Figure 3C and 3D (F: PBMCs from healthy donors, G: PBMCs from patients with SLE).

### Supplemental Fig. 4

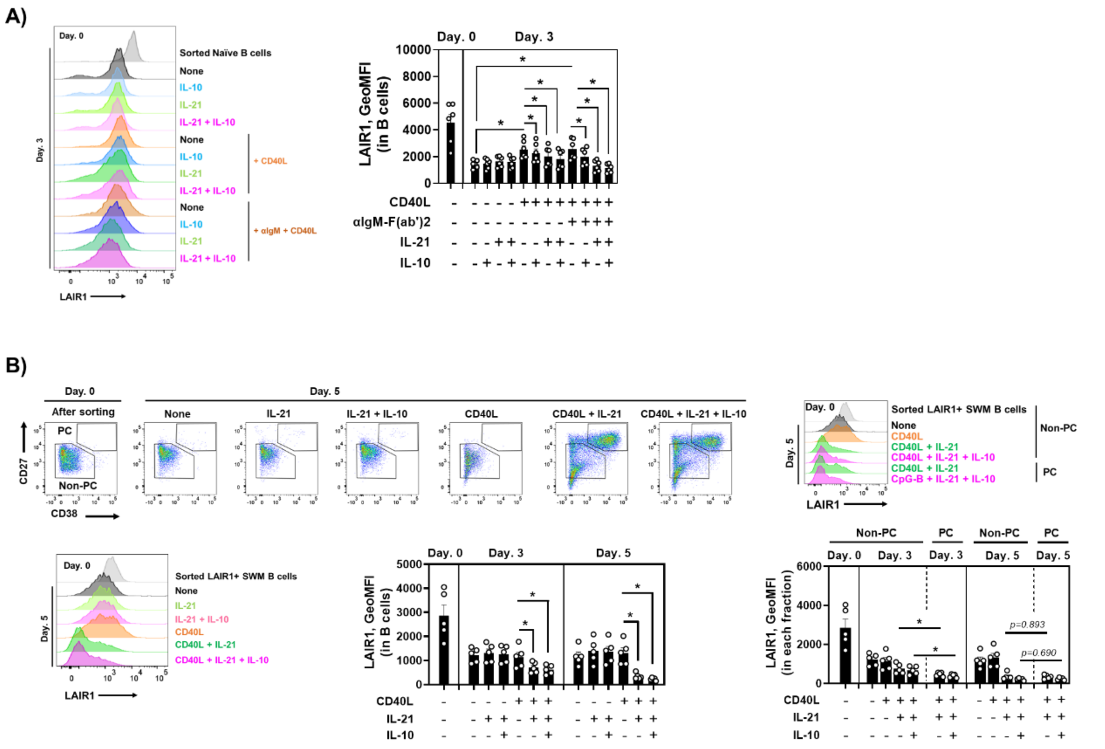

(A) The effects of IL-21 and IL-10 on LAIR1 expression in naïve B cells co-stimulated with CD40 or CD40/BCR in 3 days culture (N=6). (B) The effects of IL-21 and IL-10 on LAIR1 expression in SWM B cells co-stimulated with CD40 in 5 days culture (N=5). Data are shown as mean ± SEM with each symbol representing an individual subjects. P values were calculated with the Wilcoxon signed rank tests (A) and further adjusted for multiple comparisons using the Benjamini-Hochberg method (B). Asterisks indicate significant differences (*P < 0.05).

### Supplemental Fig. 5

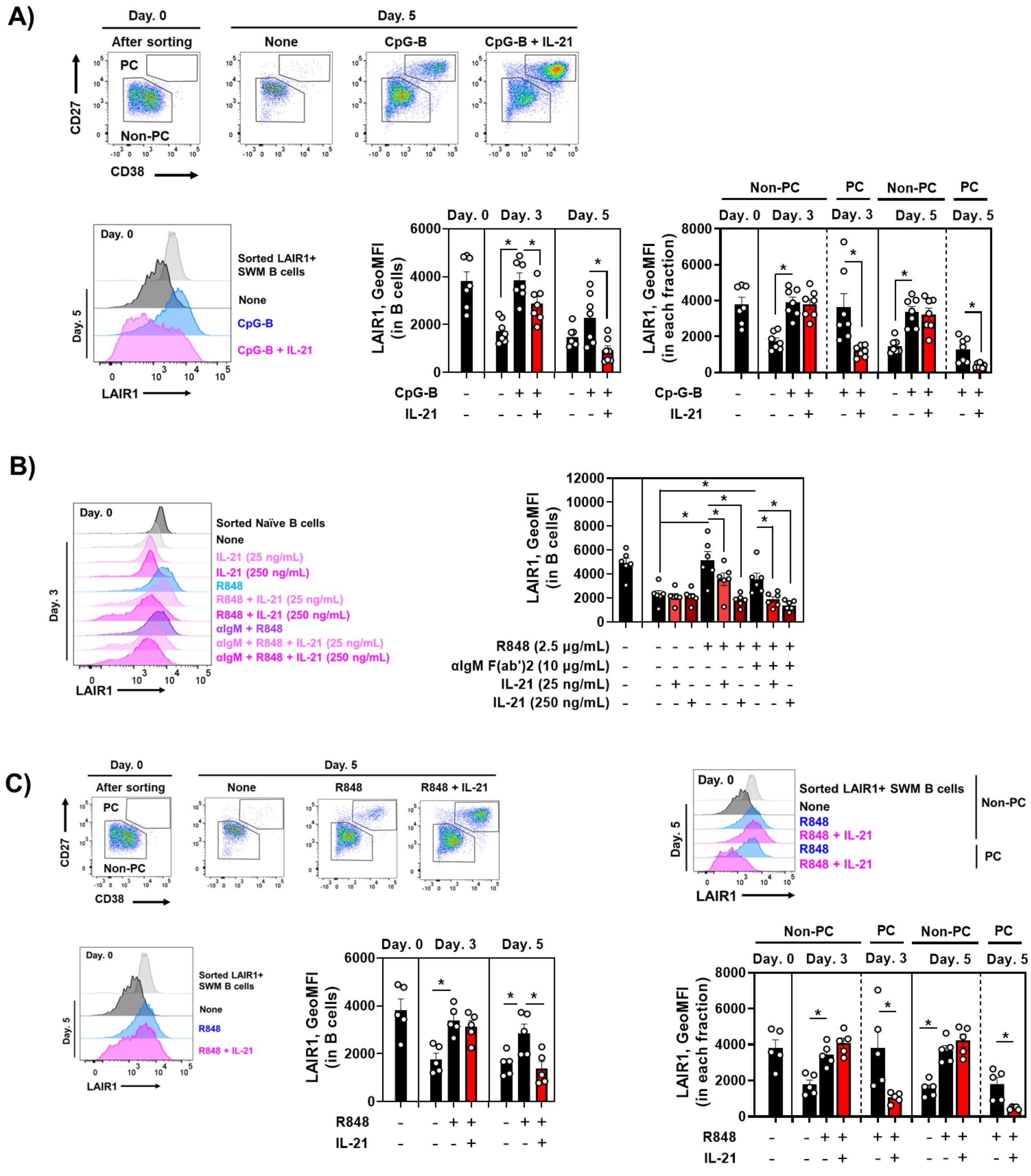

(A) The effects of IL-21 on LAIR1 expression in SWM B cells co-stimulated with TLR9 in 5 days culture (N=7). (B) The effects of IL-21 on LAIR1 expression in naïve B cells co-stimulated with TLR7 or TLR7/BCR in 3 days culture (N=6). (C) The effects of IL-21 on LAIR1 expression in SWM B cells co-stimulated with TLR7 in 5 days culture (N=5). Data are shown as mean ± SEM with each symbol representing an individual subjects. P values were calculated with the Wilcoxon signed rank tests (C) and further adjusted for multiple comparisons using the Benjamini–Hochberg method (A and B). Asterisks indicate significant differences (*$P < 0.05$).

### Supplemental Fig. 6

A)

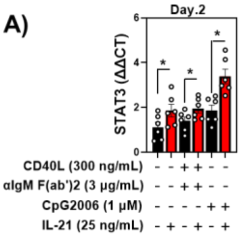


B)

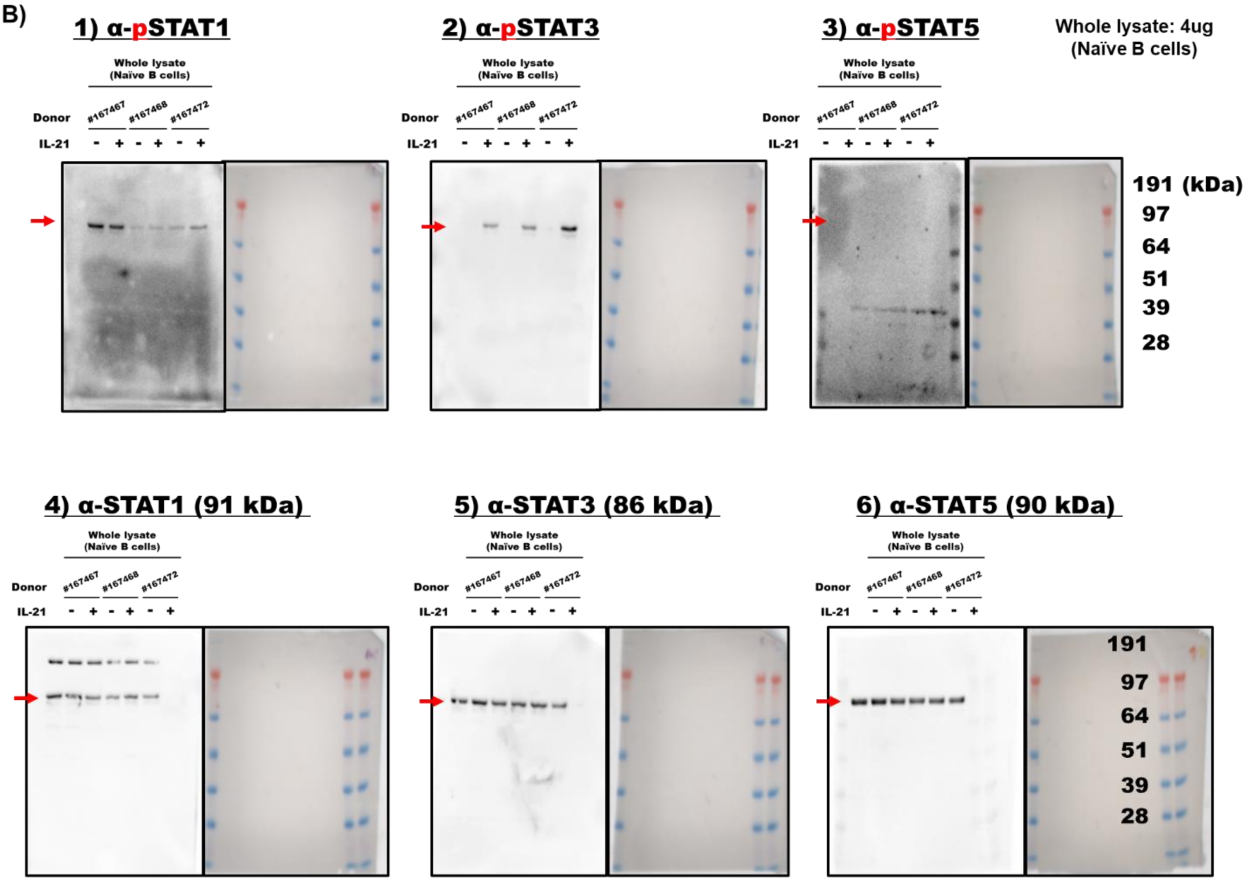

C)

### 1) α-LaminB1 (66 kDa)

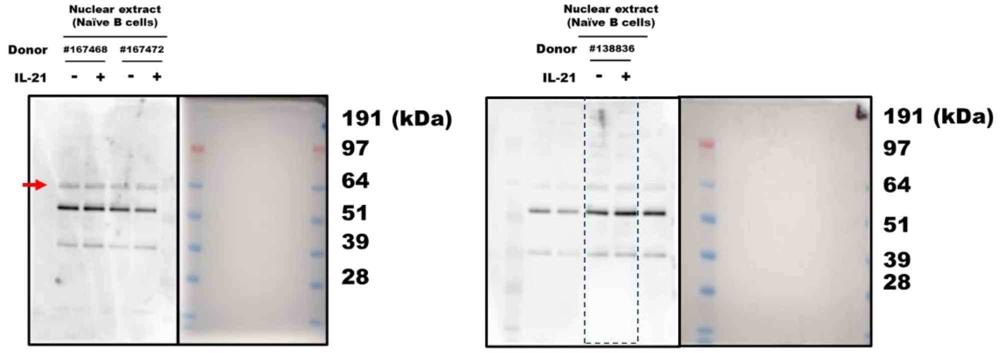


### 2) α-STAT1 (91 kDa)

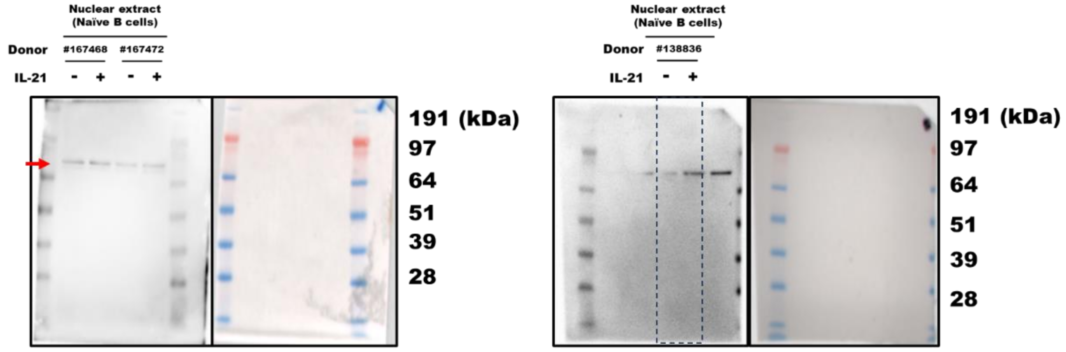


### 3) α-STAT3 (86 kDa)

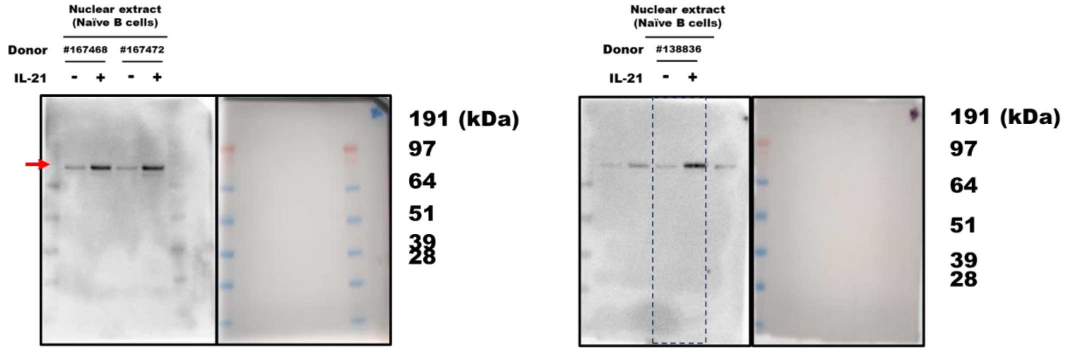


### 4) α-STAT5 (90 kDa)

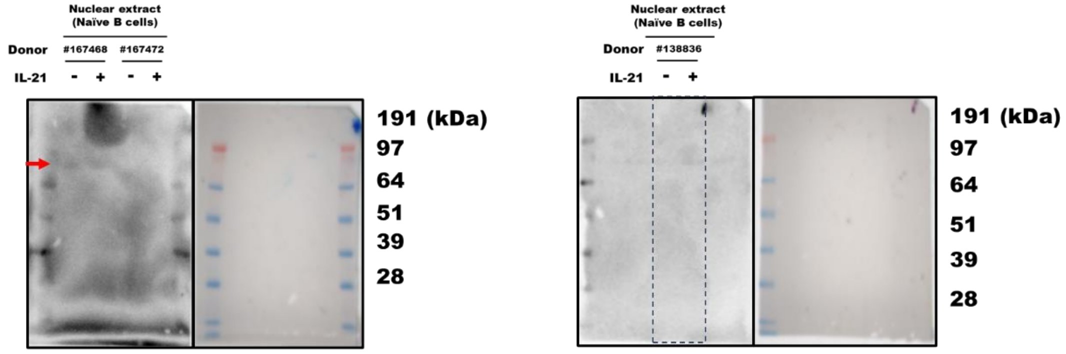

D)

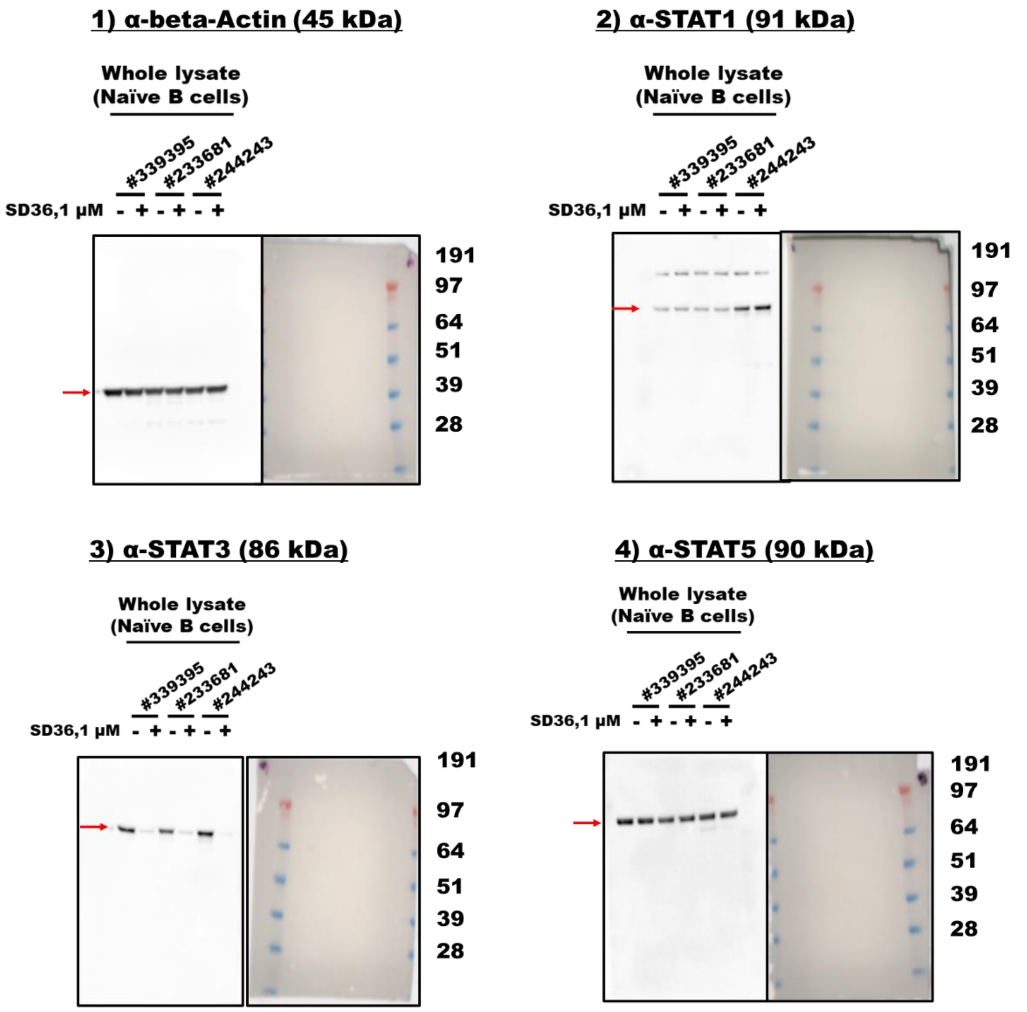
1) α-beta-Actin (45 kDa)
Whole lysate
(Naïve B cells)
#339395
#233681
#244243
SD36,1 μM - + - + - +
191
97
64
51
39
28
2) α-STAT1 (91 kDa)
Whole lysate
(Naïve B cells)
#339395
#233681
#244243
SD36,1 μM - + - + - +
191
97
64
51
39
28
3) α-STAT3 (86 kDa)
Whole lysate
(Naïve B cells)
#339395
#233681
#244243
SD36,1 μM - + - + - +
191
97
64
51
39
28
4) α-STAT5 (90 kDa)
Whole lysate
(Naïve B cells)
#339395
#233681
#244243
SD36,1 μM - + - + - +
191
97
64
51
39
28

E)

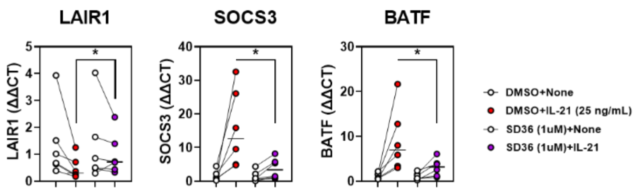
LAIR1
SOCS3
BATF
LAIR1 (ΔΔCT)
SOCS3 (ΔΔCT)
BATF (ΔΔCT)
DMSO+None
DMSO+IL-21 (25 ng/mL)
SD36 (1uM)+None
SD36 (1uM)+IL-21

(A) The effects of IL-21 on STAT3 mRNA expression in naïve B cells co-stimulated with TLR9 or CD40/BCR in 2 days culture (N=6) (B) Full images of membranes for western blotting in Figure 6C (left). (C) Full images of membranes for western blotting in Figure 6C (right). (D) Full images of membranes for western blotting in Figure 6E. (E) the effect of SD36 (1 μM) against mRNA expression of LAIR1, STAT3 and BATF induced by IL-21 (N=6). Data are shown as mean ± SEM with each symbol representing an individual subjects. P values were calculated with the Wilcoxon signed rank tests (A and E). Asterisks indicate significant differences (*P < 0.05).

**Supplemental Fig. 7**

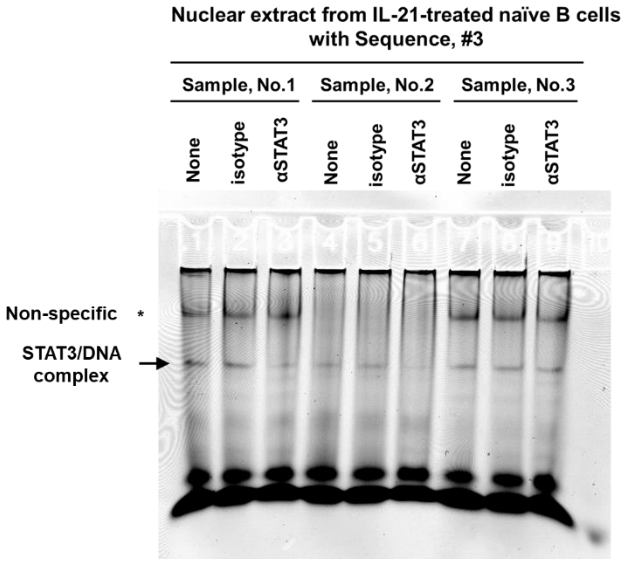


(A) Full image of EMSA of Figure 7B.

### Supplementary Table. 1 List of reagents

#### For cell isolation

| Reagent | Catalog number | supplier |
|---|---|---|
| Ficoll-Paque PLUS | 17144003 | Cytiva |
| EasySep Human CD4+ T Cell Isolation Kit | 17952 | Stem cell technologies |
| EasySep Human B Cell Isolation Kit | 17954 | Stem cell technologies |
| EasySep Human Naïve B Cell Isolation Kit | 17254 | Stem cell technologies |
| EasySep™ Human Pan-B Cell Enrichment Kit | 19554 | Stem cell technologies |

#### For cell culture and stimulation

| Reagent | Catalog number | supplier |
|---|---|---|
| X-VIVO15 Hematopoietic Serum-Free Culture Media | 04-418Q | Lonza |
| anti-IgG F(ab')2 | 2042-01 | SouthernBiotech |
| anti-IgM F(ab')2 | 109-066-129 | Jackson ImmunoResearch |
| biotinylated anti-IgG F(ab')2 | 2042-08 | SouthernBiotech |
| biotinylated anti-IgM F(ab')2 | 109-066-129 | Jackson ImmunoResearch |
| biotinylated anti-LAIR1 polyclonal antibody | BAF2664 | R&D Systems |
| biotinylated isotype antibody | BAF108 | R&D Systems |
| CpG-B (CpG2006) | tlrl-2006-1 | invivogen |
| Dynabeads Human T-Activator CD3/CD28 Beads | 11161D | Gibco |
| Recombinant Human BAFF | 310-13 | PeproTech |
| Recombinant Human IFNγ | 285-IF-100 | R&D Systems |
| Recombinant Human IL-2 | 200-02 | PeproTech |
| Recombinant Human IL-10 | 1064-ILB-010 | R&D Systems |
| Recombinant Human IL-21 | 8879-IL-010/CF | R&D Systems |
| Recombinant Human IL-21R Fc chimera protein | 991-R2-100 | R&D Systems |
| MEGA CD40L | ALX-522-110-C010 | Enzo |
| R848 (Resiquimod) | tlrl-r848-1 | invivogen |
| Streptavidin | 189730 | Sigma-Aldrich |
| SD36 | HY-129602 | MedChemExpress |
| Ultra-LEAF Purified Human IgG1 Isotype Antibody | 403502 | Biolegend |

#### For fixation and permeabilization

| Reagent | Catalog number | supplier |
|---|---|---|

| | | |
|---|---|---|
| BD Cytofix™ Fixation Buffer | 554655 | BD |
| Perm Buffer II | 558052 | BD |
| Foxp3 / Transcription Factor Staining Buffer Set | 00-5523-00 | eBioscience |

### For membrane array

| Reagent | Catalog number | supplier |
|---|---|---|
| Proteome Profiler Human Phospho-Immunoreceptor Array Kit | ARY004B | R&D Systems |

### For RNA isolation and qPCR

| Reagent | Catalog number | supplier |
|---|---|---|
| Chloroform | C2432 | Sigma-Aldrich |
| Ethanol | 2701 | Decon Labs |
| iScript cDNA Synthesis Kit | 1708891 | Bio-Rad |
| QIAzol Lysis Reagent | 79306 | QIAGEN |
| RNase-Free DNase Set | 79254 | QIAGEN |
| RNeasy Mini Kit | 74106 | QIAGEN |
| TaqMan Fast Advanced Master Mix for qPCR | 4444557 | Applied Biosystems |
| TRIzol Reagent | 15596018 | Invitrogen |

### For western blotting

| Reagent | Catalog number | supplier |
|---|---|---|
| Anti-β-Actin Antibody, polyclonal | 4967 | CST |
| Anti-Lamin B1 Antibody, polyclonal | Ab16048 | Abcam |
| Anti-STAT1 Antibody, monoclonal (D1K9Y) | 14994 | CST |
| Anti-STAT3 Antibody, monoclonal (D3Z2G) | 12640 | CST |
| Anti-STAT5 Antibody, monoclonal (D2O6Y) | 94205 | CST |
| Anti-pSTAT1 Antibody (pY701), monoclonal (58D6) | 9167 | CST |
| Anti-pSTAT3 Antibody (pY705), monoclonal (D3A7) | 9145 | CST |
| Anti-pSTAT5 Antibody (pY694), monoclonal (C71E5) | 9314 | CST |
| Anti-rabbit IgG, HRP-linked Antibody | 7074 | CST |
| BSA Fraction V | 10735108001 | Sigma-Aldrich |
| Halt Phosphatase Inhibitor Single-Use Cocktail | 78428 | Thermo Scientific |
| Halt Protease Inhibitor Cocktail (100X) | 78429 | Thermo Scientific |
| iBlot 2 Transfer Stacks, PVDF, regular size | IB24001 | Invitrogen |
| MOPS SDS Running Buffer (20X) | NP0001 | Invitrogen |

| | | |
|---|---|---|
| NE-PER Nuclear and Cytoplasmic Extraction Reagents | 78833 | Thermo Scientific |
| Non-fat dry milk (skim milk) | M0841 | Lab Scientific |
| NuPAGE™ Bis-Tris Mini Protein Gels, 4–12% | NP0323BOX | Invitrogen |
| NuPAGE LDS Sample Buffer (4X) | NP0007 | Invitrogen |
| NuPAGE MOPS SDS Running Buffer (20X) | NP0001 | Invitrogen |
| NuPAGE Sample Reducing Agent (10X) | NP0009 | Invitrogen |
| Pierce Rapid Gold BCA Protein Assay Kit | A53226 | Thermo Scientific |
| RIPA Lysis and Extraction Buffer | 89901 | Thermo Scientific |
| SeeBlue Plus2 Pre-stained Protein Standard | LC5925 | Invitrogen |
| SuperSignal West Pico PLUS Chemiluminescent Substrate | 34577 | Thermo Scientific |
| TRIS-buffered saline (TBS, 10X) pH 7.4 | J62938.K2 | Thermo Scientific Chemicals |
| Tween 20 | P9416 | Sigma-Aldrich |

### For EMSA

| Reagent | Catalog number | supplier |
|---|---|---|
| 5% Mini-PROTEAN TBE Gel | 4565015 | Bio-Rad |
| Anti-STAT3, rabbit polyclonal | 06-596 | Sigma-Aldrich |
| Isotype control, rabbit | Z25307 | Life technologies |
| Odyssey EMSA Kit | 829-07910 | LICOR bio |
| TBE Buffer, 10X Solution | BP1333-1 | Thermo Scientific |

### Supplementary Table. 2 List of antibodies for flow cytometry and sorting

| Antigen | Fluorochrome | Clone | Catalog number | supplier |
|---|---|---|---|---|
| CD3 | BV510 | UCHT1 | 300448 | Biolegend |
| CD4 | PerCP-Cy5.5 | OKT4 | 317428 | Biolegend |
| CD11c | BV421 | 3.9 | 301628 | Biolegend |
| CD14 | BV510 | M5E2 | 301842 | Biolegend |
| CD16 | BV510 | 3G8 | 302048 | Biolegend |
| CD19 | PerCP-Cy5.5 | HIB19 | 302230 | Biolegend |
| CD19 | APC | HIB19 | 302212 | Biolegend |
| CD19 | BV421 | HIB19 | 302234 | Biolegend |
| CD19 | BV510 | HIB19 | 302242 | Biolegend |
| CD20 | BV510 | 2H7 | 302340 | Biolegend |
| CD21 | FITC | Bu32 | 354909 | Biolegend |
| CD24 | BV421 | ML5 | 311122 | Biolegend |
| CD27 | BUV395 | L128 | 563815 | BD |
| CD27 | BUV395 | O323 | 751677 | BD |
| CD27 | APC | O323 | 302810 | Biolegend |
| CD27 | BV421 | O323 | 302824 | Biolegend |
| CD38 | BV711 | HIT2 | 303528 | Biolegend |
| CD45RA | FITC | HI100 | 304106 | Biolegend |
| CD45RB | FITC | MEM-55 | 310206 | Biolegend |
| CD56 | BV510 | HCD56 | 318340 | Biolegend |
| CD95 | APC | DX2 | 305612 | Biolegend |
| CD123 | BV510 | 6H6 | 306022 | Biolegend |
| CD185/CXCR5 | APC | J252D4 | 356908 | Biolegend |
| CD185/CXCR5 | BV421 | J252D4 | 356920 | Biolegend |
| CD279/PD-1 | PE | EH12.2H7 | 329906 | Biolegend |
| CD305/LAIR1 | PE | DX26 | 550811 | BD |
| CD305/LAIR1 | AF647 | NKTA255 | 342802 | Biolegend |
| IgA | APC | IS11-8E10 | 130-113-472 | Miltenyi Biotec |
| IgA | PE-Vio770 | IS11-8E10 | 130-113-477 | Miltenyi Biotec |
| IgD | FITC | IA6-2 | 348206 | Biolegend |
| IgD | BV421 | IA6-2 | 348226 | Biolegend |
| IgD | BV510 | IA6-2 | 348220 | Biolegend |

| | | | | |
|---|---|---|---|---|
| IgG | FITC | G18-145 | 555786 | BD |
| IgG | BV605 | G18-145 | 563248 | BD |
| IgM | FITC | MHM-88 | 314506 | Biolegend |
| IgM | PerCP-Cy5.5 | MHM-88 | 314512 | Biolegend |
| IgM | BV605 | MHM-88 | 314524 | Biolegend |
| pAKT (pS473) | AF488 | M89-61 | 560404 | BD |
| pERK (pT202/pY204) | AF647 | 20A | 612593 | BD |
| pSTAT1 (pY701) | BV421 | 4a | 562985 | BD |
| pSTAT3 (pY705) | AF647 | 4/P-STAT3 | 557815 | BD |
| pSTAT5 (pY694) | AF488 | 47/Stat5(pY694) | 612598 | BD |
| STAT3 | PE | M59-50 | 560391 | BD |
| IRF4 | FITC | 3E4 | 11-9858-82 | eBioscience |
| T-bet | PE | O4-46 | 561268 | BD |
| Streptavidin-APC conjugated |  | ----- | 405207 | Biolegend |
| Fc Blocking Reagent human |  | ----- | 130-059-901 | Miltenyi Biotec |
| Fixable Viability Dye eFluor™ 780 |  | ----- | 65-0865-14 | eBioscience |
| UltraComp eBeads™ Compensation Beads |  | ----- | 01-2222-42 | invitrogen |

### Supplementary Table. 3 List of Taqman Probe

| Gene name | Catalog number | supplier |
|---|---|---|
| BATF | Hs00232390_m1 | Thermo Fisher Scientific |
| BCL6 | Hs00153368_m1 | Thermo Fisher Scientific |
| EIF1b | Hs00271856_m1 | Thermo Fisher Scientific |
| IL21R | Hs00222310_m1 | Thermo Fisher Scientific |
| IRF4 | Hs00180031_m1 | Thermo Fisher Scientific |
| LAIR1 | Hs00253790_m1 | Thermo Fisher Scientific |
| PRDM1 | Hs00153357_m1 | Thermo Fisher Scientific |
| SOCS3 | Hs02330328_s1 | Thermo Fisher Scientific |
| STAT3 | Hs00374280_m1 | Thermo Fisher Scientific |

### Supplementary Table. 4 List of EMSA Probe

| Sequence # | Conjugated | promoter sequence | supplier |
|---|---|---|---|
| #1: -1442 to -1467 bp | IRDye800 | GCTCTAGTTCCAAGTAAGATCCTTAT | Integrated DNA Technologies |
| #2: -731 to -755 bp | IRDye800 | CCTCAACCTTCCAAGTAGCTGTGCCC | Integrated DNA Technologies |
| #3: -69 to -94 bp | IRDye800 | GCGACATTTTTCAGAAACCAAGGCCC | Integrated DNA Technologies |

### Supplementary Table. 5 Clinical characteristics of patients with SLE

| | SLE (N=9) | Healthy Donor (N=9) |
|---|---|---|
| Age | 43.6 (27-61) | 37.2 (16-64) |
| Sex (N=female/male) | 8/1 (female: 88.9%) | 9/0 (female: 100%) |
| SLEDAI | 2.2 (0-8) | ----- |
| Clinical symptoms | Renal: 0 (0%)<br>Arthritis: 1 (11.1%)<br>Rash: 1 (11.1%) | ----- |
| Current medications | Prednisolone: 4 (44.4%)<br>MMF: 3 (33.3%)<br>HCQ: 6 (66.7%)<br>Belimumab: 1 (11.1%)<br>Anifrolumab: 1 (11.1%)<br>Colchicine: 1 (11.1%) | ----- |
| Anti-dsDNA IgG<br>%positive (12 IU/mL~) | <br>7 (77.8%), positive: 16 - >1000 IU/mL | ----- |
| Complement<br>C3 (mg/dL)<br>C4 (mg/dL) | <br>121.9 (102-166)<br>22.4 (10-40) | ----- |

### Supplementary Table. 6 Terminology and B cell definition

<table>
<tr><td>Terminology</td><td>Primary population</td><td>Secondary population</td></tr>
<tr><td>Naïve</td><td rowspan="3">Naïve<br>: CD19$^{+}$ CD27$^{-}$ IgD$^{+}$</td><td>---</td></tr>
<tr><td>Early memory (CD45RB$^{+}$ naive)</td><td>CD45RB$^{+}$ CD38$^{lo/+}$</td></tr>
<tr><td>Activated naïve</td><td>CD21$^{-}$ CXCR5$^{-}$</td></tr>
<tr><td>DN1</td><td rowspan="3">DN<br>: CD19$^{+}$ CD27$^{-}$ IgD$^{-}$</td><td>CD21$^{+}$ CXCR5$^{+}$</td></tr>
<tr><td>DN2</td><td>CD21$^{-}$ CXCR5$^{-}$</td></tr>
<tr><td>ABC-like (in vitro)</td><td>CD11c$^{+}$ Tbet$^{+}$</td></tr>
<tr><td>USWM</td><td>USWM: CD19$^{+}$ CD27$^{+}$ IgD$^{+}$</td><td>---</td></tr>
<tr><td>SWM</td><td>SWM: CD19$^{+}$ CD27$^{+}$ IgD$^{-}$</td><td>---</td></tr>
</table>
